# Epithelial Plasticity and an Immune Suppressive Microenvironment Underpin Tumour Budding in Colorectal Cancer

**DOI:** 10.1101/2025.03.05.641608

**Authors:** Phimmada Hatthakarnkul, Natalie Fisher, Leonor Schubert Santana, Holly Leslie, Assya Legrini, Aula Ammar, Amna Ahmed Mohemmed Malty, Kathryn A. F. Pennel, Ian Powley, Courtney Bull, Peter G. Alexander, Janos Sztanko, Jean A. Quinn, Jennifer Hay, Hannah Morgan, Claire Kennedy Dietrich, Yoana Doncheva, Gerard Lynch, Noori Maka, Hester C. van Wyk, Andrew D. Campbell, Xiao Fu, Donald C. McMillan, Owen J. Sansom, Philip Dunne, Nigel Jamieson, Campbell Roxburgh, Chanitra Thuwajit, Joanne Edwards

## Abstract

**Background:** Tumour budding (TB), defined as a small cluster of up to four cells at the invasive front of the tumour, is a well-established independent and robust prognostic biomarker in colorectal cancer (CRC). This is strongly associated with adverse clinicopathological features and poor survival outcomes. Despite its clinical relevance, the precise underlying mechanism responsible for TB phenomenon remains unclear.

**Methods:** Multi-omic approaches from bulk, regional GeoMx and Spatial Molecular Imager (SMI) RNA were used to identify the underlying mechanism of TB and its possible correlation with tumour microenvironment (TME) in CRC tissue. The results were validated using immunohistochemistry (IHC) and multiplex immunofluorescence (mIF) staining.

**Results:** Patients with high TB experience worse outcomes and associate with adverse clinical factors across two independent CRC cohorts. Bulk and regional RNA expression analyses reveal that tumours with high TB are significantly enriched for TNF-α and TGF-β signatures in both cohorts. Single cell CosMx SMI analysis confirmed TB cells exhibit higher expression of these signatures than adjacent invasive edge tumour cells. Elevated cyclinD1 expression was also observed within TB, and high cyclinD1 levels tend to experience poorer CRC prognosis. Furthermore, regional bulk RNA expression within the non-tumour (PanCK-) invasive edge areas demonstrated that tumours exhibiting high TB revealed the significantly differential expressions of immune-related genes (e.g. *CD3*, *NKG7*, *IL6*, *CXCR6*, *CD47*, *IFNAR1* and *VSIR*). Single cell CosMx SMI analysis revealed that cancer-associated fibroblasts (CAFs) were physically the closest cells to TB cells. This spatial proximity was confirmed at the protein level using mIF, where the distance from TB to CD68+ macrophages predicted significantly poorer CRC outcomes.

**Conclusion:** This multi-omic study confirms the prognostic significance of TB in CRC patients across two independent cohorts. Our findings highlight that TNF-α and TGF-β signalling play a crucial role in budding cells development by regulating cyclinD1. Furthermore, the transcriptomic analysis reveals an immunosuppressive niche characterised by reduced immune activity and close spatial interactions with CAFs and macrophages Ultimately, this study provides valuable insight into TB’s underlying mechanism and its complex interactions within the TME. This could provide a foundation for developing targeted therapeutic strategies in CRC.

## Background

Colorectal cancer (CRC) is the third most diagnosed cancer and the second most lethal malignancy worldwide (2). Incidence rates of CRC have been decreasing due to the development of screening programs (3). However, 30% of CRC patients develop synchronous or metachronous metastasis (4). Of these, fewer than 20% with metastatic CRC are alive 5 years after diagnosis (5). Thus, one of the main challenges in cancer research is to develop targeted treatments to prevent metastasis and subsequently increase the survival rate of those with CRC (6).

Tumour budding (TB) is a single cell or small cluster of up to 4 cells found at the invasive front of tumours (7) and has been consistently reported as a poor prognostic factor associated with tumour recurrence and metastases across solid tumours (8–10). TB has been reported as an independent prognostic marker associated with adverse clinicopathological factors such as lymphatic invasion, venous invasion, and disease recurrence (11–13). Recently, using the International Tumor Budding Consensus Conference (ITBCC) criteria for TB assessment, the prognostic and predictive values of TB have been reported in stage II CRC from both SACURA (14) and QUASAR (15) trials.

Although the potential prognostic value of TB in CRC has been widely reported, the molecular mechanisms underpinning TB are poorly understood (10). Recently, we published a literature review detailing the potential underlying mechanisms of TB in CRC as well as a systematic review of how TB may be associated with common CRC mutations (16, 17). Additionally, the role of inflammation in CRC has been repeatedly reported as a potential prognostic biomarker in CRC (18–20). Studies showed that a high number of TBs appear to compromise immune activity leading to invasion and metastasis in CRC (16, 21). It is essential to elucidate the molecular mechanisms driving TB, as these may represent novel therapeutic targets in CRC. To address this, we applied an integrated multi-omic strategy combining bulk transcriptomics, regional digital spatial profiling (DSP), and single-cell spatial molecular imaging (SMI) RNA analysis, complemented by proteomic validation. This comprehensive approach enables a deeper understanding of TB biology and its interplay with the tumour microenvironment (TME), providing critical insights into pathways that underpin invasion, immune evasion, and disease progression in CRC.

## Methods and Materials

### Patient Cohort

#### Discovery cohort

A cohort of stage I-III CRC patients (n=787) who underwent resection for primary operable CRC in the Glasgow Royal Infirmary (GRI, Glasgow, UK) between 1997 and 2013 formed the discovery cohort. Patients who died within 30 days of surgery, had emergency operation, or received neoadjuvant therapy were excluded. Tumour staging was carried out using 5th Edition of the AJCC/UICC-TNM staging system. Clinicopathological data were collected with a minimum of 10 years follow-up years post-resection. The study was approved by the West of Scotland Research Ethics Committee (REC 4: 22/ws/0207) and data are stored in the Greater Glasgow and Clyde Safehaven (SH21ON012).

#### Validation cohort

A publicly available cohort of stage II-III Colon Cancer (CC) patients (n=85), who underwent curative surgery at the Department of Surgery, National Defense Medical College Hospital (Saitama, Japan) between 2002-2003 was used as the validation cohort. The cohort was downloaded from GEO under accession number GSE143985 (1). The data was obtained and processed by NF and CB.

### Tumour budding assessment

Following the guidelines from ITBCC 2016 (7), budding scores were assessed in both discovery and validation cohorts. Full CRC sections were stained with Haematoxylin and Eosin (H&E). Ten specific areas (0.785 mm^2^) at the invasive tumour front were marked and the number of TBs were determined at 20x magnification. Budding status in the discovery cohort was determined by HW and PH. In validation cohort, TB status was previously assessed, and data was obtained and processed by NF. The densest area of buds represented the budding profile of each CRC case. Low and high TB phenotypes were based on the number of TB (<10 as low and ≥10 as high) found at the invasive area (Figure 1A).

**Figure 1.**
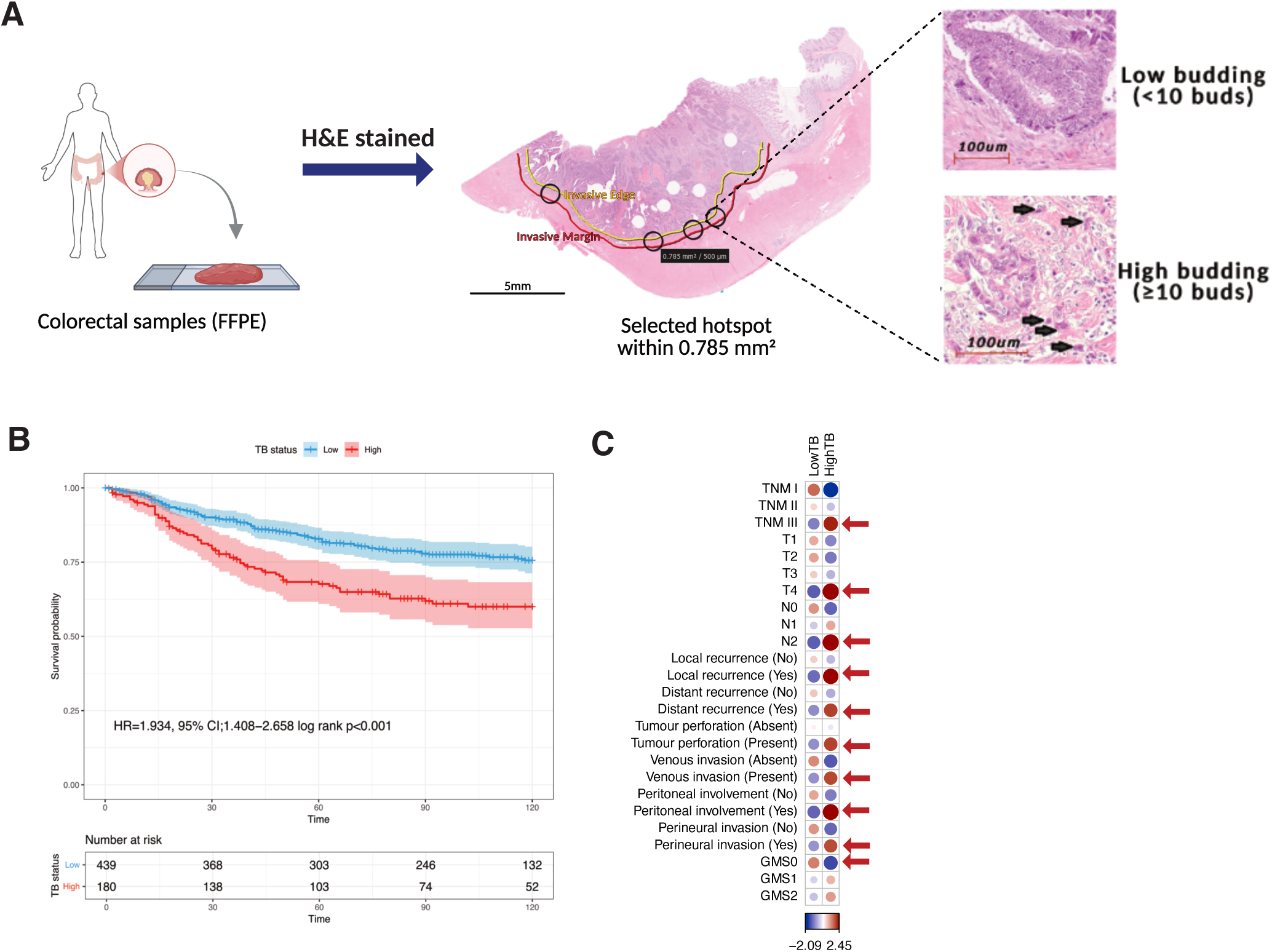
(A) Colorectal cancer tissue sample was stained with H&E the invasive edge was identifed within 0.785mm^2^ area from to the tumour invasive edge to define budding phenotype (low <10 TB and high >10 TB). (B) Kaplan-Meier survival analysis based on tumour budding phenotype for cancer specific survival (CSS) in discovery cohort. (C) The correlation plot for the Pearson’s chi square test showed the residuals for TB phenotypes against clinical factors; positive residuals are in blue, suggesting a positive association and negative residuals are red, suggesting a negative association. The bigger size of the circle, the more significant association was found, red arrow indicates the statistically significant association.

### Bulk Whole Transcriptomic Assay (TempO-Seq^TM^)

Of the 787 patients from discovery cohort, 764 single 4µm thickness formalin-fixed paraffin-embedded (FFPE) from CRC resections were used for Templated Oligo-Sequencing (22, 23). (See Supplementary Method 1)

### Regional bulk transcriptomic (GeoMx Digital Spatial Profiler; DSP)

Whole CRC tissue sections from the discovery cohort were selected (n=12, 6 low and 6 high TB) and GeoMx RNA Immune-Oncology Pathways panel (84 targets) used to identify the gene expression within regions of interest (ROIs) proximal and distal to the invasive front of either tumour-rich (PanCK+) or non-tumour (PanCK-) regions (i.e. stroma-rich region) (Figure 2A). Additionally, a tissue microarray (TMA), of tumour invasive front from the discovery cohort with low (n=20) and high (n=23) TB samples underwent human whole transcriptomic atlas (WTA) GeoMx RNA panels (18,677 targets) analysis (Supplementary Figure S6A). Target gene list from both panels are shown in Supplementary Table S1. The DSP workflow is described in Supplementary Method 2.

**Figure 2.**
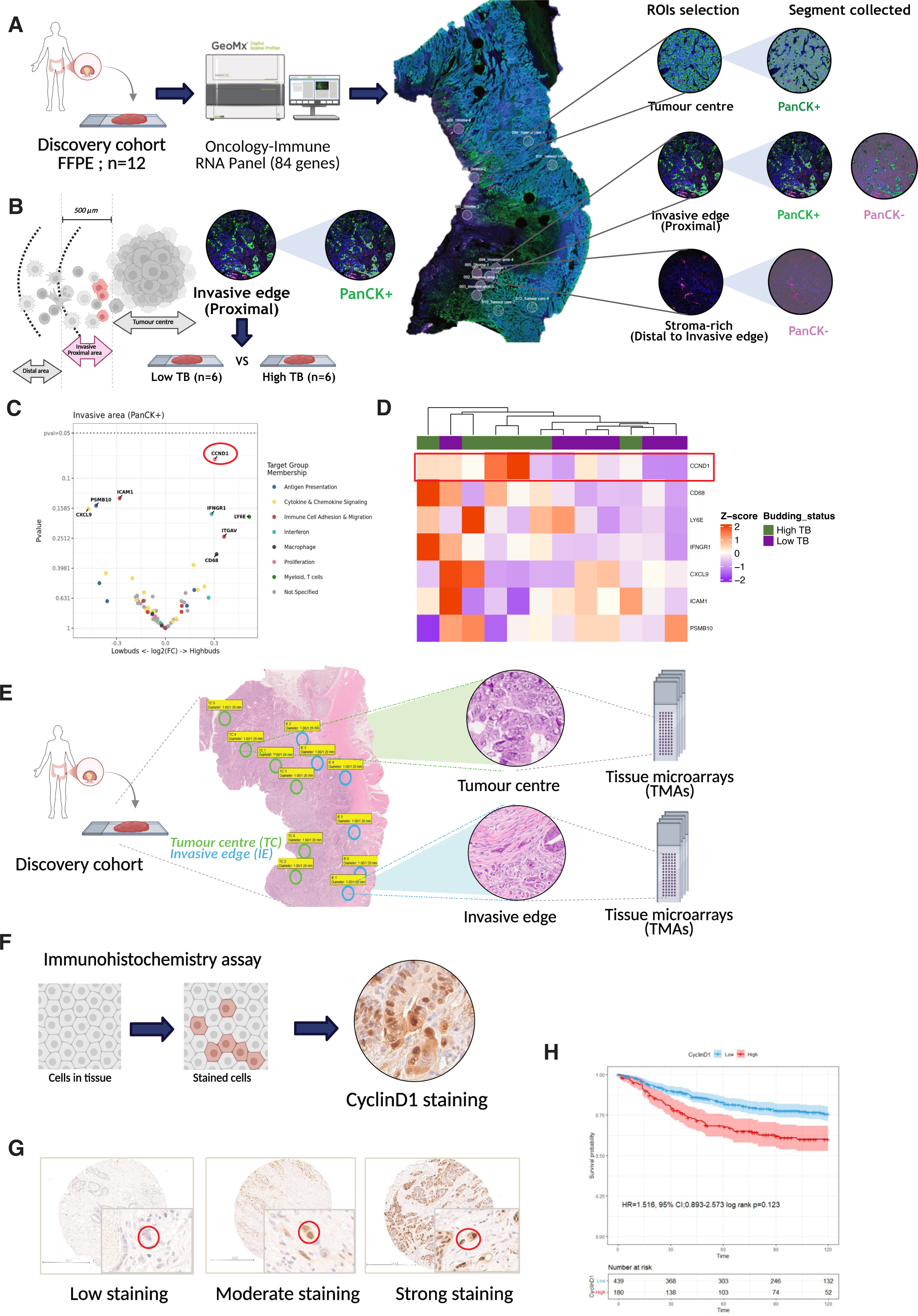
(A) The illustration shows the workflow of the GeoMx RNA Immune-Oncology panel: regions of interest were selected (tumour centre, invasive edge, and stroma-rich areas), and segmentation was collected based on PanCK staining (epithelial; PanCK+, non-epithelial; PanCK-). (B) Region of tumour invasive area (PanCK+) used to determine the differential genes expression between tumours with low and high TB. (C) Volcano plot of gene expression in the tumour invasive area (PanCK+) between tumours with high and low budding profile (n=12; low 6 and high 6). (D) Heatmap illustrating the expression of the genes in the two different groups. (E) The graphic shows the area selected for tissue microarray construction (tumour centre and invasive edge) from the discovery cohort. (F) IHC staining of CyclinD1 protein expression nuclear staining in CRC invasive edge TMA (created in BioRender Pennel, K. (2026) https://BioRender.com/36d02i4) (G) CyclinD1 staining shows the different expression of cyclinD1 when assessed within TB cells. (H) Kaplan-Meier survival analysis based on cyclinD1 expression for cancer specific survival (CSS) in TB cells.

### Immunohistochemistry (IHC)

IHC was performed on a TMA previously made from both tumour centre and invasive front of the discovery cohort (Figure 2E). The weighted histoscore was used to assess nuclear cyclinD1 protein expression as explained in Supplementary Method 3.

### CosMx Spatial Molecular Imager (SMI) – Single-cell spatial transcriptomics

CosMx SMI technology deploying the human 6K discovery RNA Panel (6,175 genes, including 20 negative control probes) was performed in 210 Fields of View (FOVs) from TMA cores from the discovery cohort (tumour centre; n=8, and invasive edge; n=4) (See Supplementary Method 4). 8 FOVs from invasive edge were selected (Supplementary Figure S1). Target gene list from human 6K panel is shown in Supplementary Table S2. Data analysis was performed by LS as described in Supplementary Method 5. The workflow is illustrated in Supplementary Figure S8.

### Lunaphore COMET^TM^ sequential immunofluorescence (seqIF™)

seqIF™ protocol was performed using the SPYRE^TM^ Immuno-Oncology Core Panel using the COMET™ platform (Lunaphore Technologies) (See Supplementary Method 6). The OME-TIFF output contains images of DAPI for nuclear staining, two auto-fluorescence channels for background subtraction, and the 16 single marker channels which were used for downstream image analysis. HORIZON^TM^ software (*v2.4.0.0)* was used for background subtraction. Subtracted images were then imported to Visiopharm (*v2025.02.2.18022*) to quantify cell density and conduct further spatial analysis (See Supplementary Method 7).

### Statistical analysis

In the discovery cohort, survival analysis was conducted using the *survival* (*v3.5-3*) R packages. Pearson’s χ2 test assessed the relationship between TB status and clinicopathological features. The likelihood ratio test was used when required. Patient survival was assessed by Kaplan-Meier analysis and log rank to test the significance. Univariate and multivariate Cox hazard regression was performed to estimate the hazard ratio (HR) for cancer-specific survival (CSS) and disease-free survival (DFS) to identify any significant prognostic factors in CRC patients. CSS was defined as patient survival in months from the date of surgery until recurrence or cancer-caused mortality. DFS was defined as patient survival in months without disease measured from treatment completion to recurrence or death from any cause. *maxstat (v0.7-25)* and *survminer (v0.4.9)* packages were utilised to determine optimal thresholds for weighted histoscores and counts from IHC and mIF staining.

In the validation cohort, disease free survival analysis was conducted using the *survminer* (*v0.4.9*) and *survival* (*v3.5-3*) R packages. Univariate and multivariate Cox hazard regression was performed to estimate the hazard ratio (HR) for DFS. This was calculated for budding status with the *MASS* (*v7.3-58.2*) and *finalfi*t (*v1.0.8*) R packages. Correlation analysis was conducted in the *stats* (*v4.2.3*) and visualised within the *corrplot* R package (*v0.92*).

Two-sided *t*-tests were used to compare between two groups and ANOVA with Welch’s correction for 3+ group comparison. In this study, a P-value or P.adj of less than 0.05 was considered to be statistically significant.

## Results

### Clinicopathological parameter of colorectal cancer patient’s cohort

In the discovery cohort, after exclusion criteria applied, 644 patients were included in the analysis. 205 (32%) patients were under 65, 208 (32%) were between 65-74 and 231 (36%) were over 75 years of age, with 353 males and 291 females. 471 (73%) and 173 (27%) patients were diagnosed with colon and rectal cancer respectively. Cancer-specific survival (CSS) and DFS were both employed as primary endpoints for patient survival. The clinical characteristics of the discovery cohort are detailed in Supplementary Table S4. Of these 644 cases, TB could not be identified in 25 cases as the image quality was inadequate (i.e. staining quality too faint), therefore 619 patients were included in the final analysis in the discovery cohort.

The validation cohort was comprised of 85 CC patients with DFS employed as a primary endpoint for patients’ survival. The mean age at diagnosis was 66.5 years. Clinical characteristics for the discovery cohort are recorded in Supplementary Table S5.

### The prognostic role of TB in CRC patients

When the discovery cohort was dichotomised into low and high TB groups, patients with a high TB phenotype (n=180) had significantly shorter CSS than those with low TB (n=439) (HR=1.934, 95% CI; 1.408-2.658; p<0.001; log-rank test) (Figure 1B). Similarly, patients with high TB (n=431) showed shorter DFS than those with low TB (n=179) (HR=1.754, 95% CI; 1.301-2.364; p<0.001; log-rank test) (Supplementary Figure S3A).

Pearson’s chi-square test revealed a strong association between high TB and adverse clinicopathological features, including TNM (TNMIII, p=0.003), T stage (T4, p=0.004), N stage (N2, p=0.002), local recurrence (Yes, p=0.006), distant recurrence (Yes, p=0.028), tumour perforation (positive, p=0.046), venous invasion (positive, p=0.006), peritoneal involvement (positive, p=0.002), perineural invasion (positive, p=0.012), and a negative association with local inflammatory scores such as Glasgow microenvironment (GMS) (24), a combined score between Klintrup–Mäkinen (KM) and tumour stromal percentage (TSP) (GMS0, p=0.046). (Figure 1C) (Supplementary Table S6).

Stratified analyses confirmed that high TB predicted shorter CSS in left-sided colon; p=0.007 and rectal; p=0.001, MMR status (pMMR; p<0.001), later disease staged (TNM III; p=0.010), negative margin involvement; p=0.001, positive venous invasion; p<0.001, negative local recurrence; p=0.002. Additionally, tumours with high TB consistently predicted the worst outcome in distance recurrence, systemic and local inflammation, regardless of clinical subgroups being analyses (Supplementary Figure S2). When analysed by tumour site, high TB was associated with the poorest CSS and DFS in both colon and rectal cancers, confirming its strong prognostic value (Supplementary Figure S3B-C). Crucially, multivariate analysis was performed and revealed that TNM stage, margin involvement, perineural invasion and TB status were independent prognostic factors for CRC CSS survival (p<0.001, p=0.007, p=0.003 and p=0.014 respectively) (Table 1).

**Table 1.**
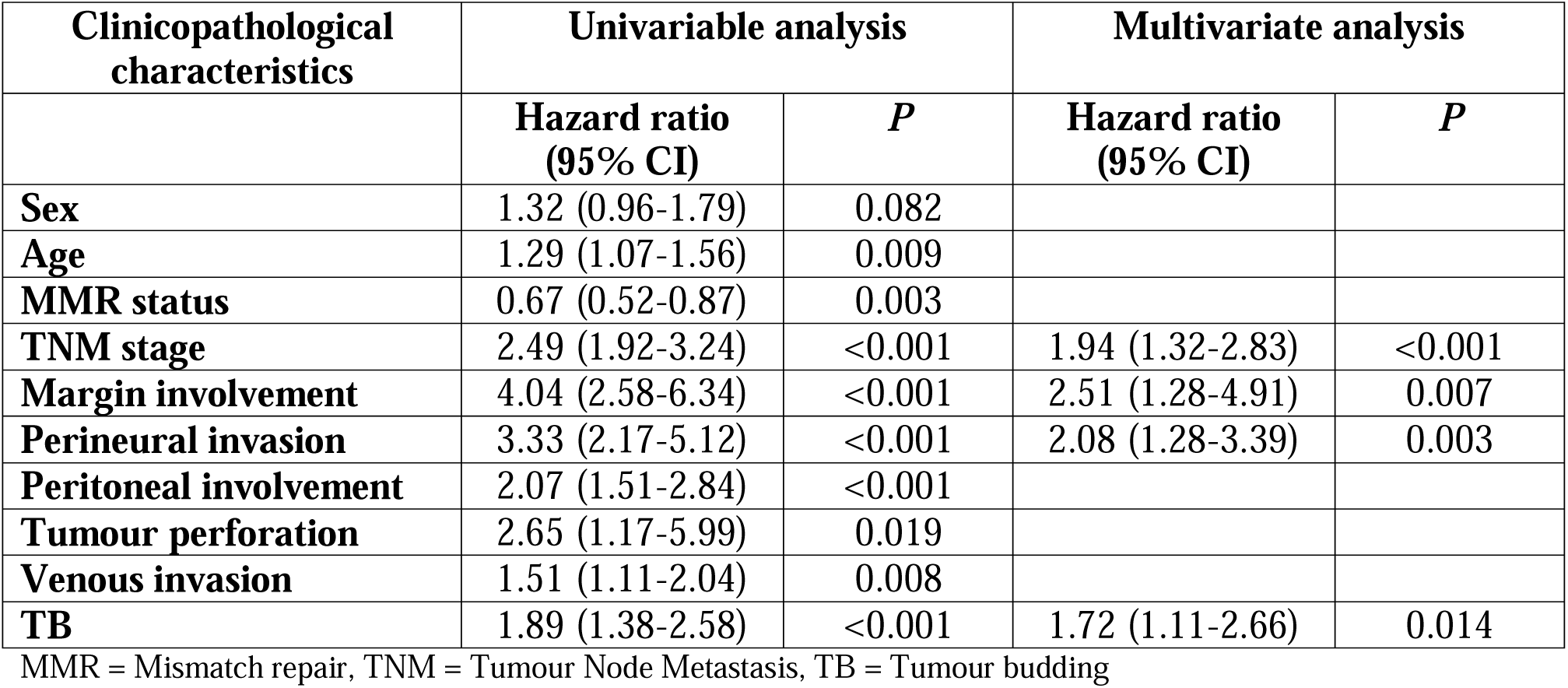
Univariate and Multivariate analysis for cancer specific survival (CSS) in discovery cohort.

| Clinicopathological characteristics | Univariable analysis |  | Multivariate analysis |  |
| --- | --- | --- | --- | --- |
|  | Hazard ratio (95% CI) | P | Hazard ratio (95% CI) | P |
| **Sex** | 1.32 (0.96-1.79) | 0.082 |  |  |
| **Age** | 1.29 (1.07-1.56) | 0.009 |  |  |
| **MMR status** | 0.67 (0.52-0.87) | 0.003 |  |  |
| **TNM stage** | 2.49 (1.92-3.24) | <0.001 | 1.94 (1.32-2.83) | <0.001 |
| **Margin involvement** | 4.04 (2.58-6.34) | <0.001 | 2.51 (1.28-4.91) | 0.007 |
| **Perineural invasion** | 3.33 (2.17-5.12) | <0.001 | 2.08 (1.28-3.39) | 0.003 |
| **Peritoneal involvement** | 2.07 (1.51-2.84) | <0.001 |  |  |
| **Tumour perforation** | 2.65 (1.17-5.99) | 0.019 |  |  |
| **Venous invasion** | 1.51 (1.11-2.04) | 0.008 |  |  |
| **TB** | 1.89 (1.38-2.58) | <0.001 | 1.72 (1.11-2.66) | 0.014 |

MMR = Mismatch repair, TNM = Tumour Node Metastasis, TB = Tumour budding

Consistent with the discovery cohort, patients with high TB (n=26) in the validation cohort also had shorter DFS outcomes compared to those with low TB (n=59) (HR=11.77, 95% CI;2.54-54.51, p<0.001) (Supplementary Figure S4A). Life tables demonstrated that 65% of patients with high budding versus 96% of patients with low budding had disease free survival 10 years after initial diagnosis. There is a significant positive correlation between high TB and clinical characteristics, using Pearson chi-square test, such as higher TNM stage (stage III, p=0.002) (Supplementary Figure S4B) (Supplementary Table S7). This is consistent with the discovery cohort as higher stage is significantly associated with high TB. Moreover, when stratified based on adverse clinical factors, patients with high TB were shown to have poorer survival such as those with stage III CRC (p=0.0011) and MSS tumours (p<0.001) (Supplementary Figure S4C-D). In concordance with the discovery cohort, TB was an independent prognostic factor for DFS in Cox regression analysis (Supplementary Table S8).

### Bulk RNA transcriptomic analysis revealed a metastatic cascade involved in tumours with high TB phenotype

To determine the biological differences between tumours with low and high TB, bulk transcriptomic analysis was performed for the discovery cohort (n=318) (Supplementary Figure S5A) and validation (n=85) (Supplementary Figure S5F) cohorts. In the discovery cohort, 9 cases were excluded as budding status was not reported, leaving 309 cases for analysis. Using the hallmark msigdb database (H), tumours with high TB were enriched for genes related to epithelial–mesenchymal transition (EMT) (padj <0.001, NES=2.46 and padj <0.001, NES=3.06), transforming growth factor beta (TGF-β) signalling (padj =0.006, NES =1.77 and padj =0.056 NES =1.47), tumour necrosis factor alpha (TNF-α) signalling via NF-κB (padj = 0.01, NES=1.50 and padj <0.001 NES=1.64) in both the discovery (Supplementary Figure S5B-C and validation cohorts respectively (Supplementary Figure S5G-H). In addition, when using a curated gene set (C2), representing canonical pathway and sets corresponding to chemical and genetic perturbation (25), it was found that tumours with high TB were enriched with genes related to advanced gastric cancer (padj <0.001, NES =3.24 and padj<0.001 NES =2.92), invasiveness (padj <0.001, NES = 3.05 and padj<0.001 NES= 3.06), up-regulated of TGF-β/EMT (padj <0.001, NES = 3.00 and padj<0.001 NES= 2.73) signatures for discovery (Supplementary Figure S5D-E) and validation cohort respectively (Supplementary Figure S5I-J).

### Expression of EMT-related genes in budding cells

To better determine epithelial gene expression, regional bulk transcriptomics GeoMx WTA analysis was performed in CRC TMAs derived from the invasive edge. Tumours with low (n=20) and high (n=23) budding regions were selected from the discovery cohort (Supplementary Figure S6A). PanCK staining was used to mask tumour epithelium (PanCK+) areas for further analysis (Supplementary Figure S6B). Interestingly, pairwise GSEA of the invasive tumour areas with high TB compared to low TB revealed an upregulation of EMT-related genes (padj <0.001, NES=1.76) and down-regulation of genes related to activation of KRAS (padj <0.001, NES=2.19) (Supplementary Figure S6C-D).

Both bulk and regional RNA expression analyses consistently demonstrated that tumours with high TB were strongly enriched for an EMT signature. Therefore, a further study was performed to investigate if TBs exhibit EMT. Immunofluorescent staining of two known EMT markers (E-cadherin and β-catenin) was performed in the invasive edge TMAs. High TB cases were selected to determine if TB were undergoing EMT (n=180) (Supplementary figure S6E). Interestingly, within the TB population, only a small proportion of TBs were found to have low E-cadherin and high β-catenin expression which represents the activation of EMT. (Supplementary figure S6F). This observation suggests that EMT may not be the sole signalling pathway driving TB in CRC, highlighting the potential involvement of alternative mechanisms.

### CyclinD1 overexpression at the invasive edge and its potential role in tumour budding

To refine our understanding of the molecular mechanisms of the budding phenotype, we examined gene expression at the tumour invasive front using regional bulk transcriptomic GeoMx DSP (Figure 2A). Using FFPE CRC full sections, tumour epithelial (PanCK+) invasive edge areas were selected to compare gene expression between cases with a low (n=6) or high (n=6) budding phenotype (Figure 2B). Although there were no statistically significant differences between the groups, *CCND1*—encoding cyclinD1, a key regulator of cell-cycle progression—was notably overexpressed in high TB cases compared to low TB (Figure 2C-D).

To increase the sample size and therefore statistical power of our analysis, TMAs constructed from cores taken at both the tumour centre and invasive edge were utilised (Figure 2E). To investigate the role of cyclinD1 and its association with TB phenotype, TMA tissue sections were stained for cyclinD1 protein using immunohistochemistry (Figure 2F). There was no difference between the weighted histoscores for cyclinD1 between low and high budding groups for both the tumour centre (Supplementary Figure S7A) and invasive edge area (Supplementary Figure S7B). However, of greater interest was the observation that increased expression of cyclinD1 within TB as assessed by weighted histoscore was associated with outcome (Figure 2G). Patients were dichotomised into those with high or low cyclinD1 within their TB, patients with high cyclinD1 (n=61) had shorter survival when compared to those with low cyclinD1 (n=88) (HR=1.516, 95% CI;0.893-2.573, p=0.123) (Figure 2H). Although this difference did not reach statistical significance, the trend suggests a potential association between elevated cyclinD1 in budding cells and poorer outcomes, warranting further investigation in larger cohorts.

### Single cell imaging revealed the possible role of NF-κB and TGF-β signalling within budding cells in CRC

To identify which biological mechanism and signalling pathway are active within the individual TB in CRC. Single cell CosMx SMI analysis was employed to pinpoint the exact molecular profile of TB in CRC. After the initial quality control and batch correction, 8 FOVs were studied focusing on areas at the invasive edge with different densities of TB. Comprehensive patient details are available and can be found in Supplementary Table S8.

Tumour cells classification was performed using the semi-supervised cell typing method InSituType (26), UMAP analysis revealed seven different tumour epithelial cell clusters (Figure 3A). Interestingly, the majority of the TB displayed an inflammatory phenotype based on their gene signature (Supplementary Figure S9). Therefore, further analyses were undertaken to assess the activation of signalling pathways in TB relative to other tumour epithelial clusters. Overall, the results confirm our hypothesis that the molecular pathways associated with TB are distinct from those in other types of tumour clusters (Supplementary Figure S10). The striking activation of NF-κB, TGF-β, cell adhesion and motility, type-I-interferon and MHC-Class I signalling pathway in TB compared to other clusters implies a specific and complex biological profile associated with increased tumour invasion, immune evasion and poor prognosis. Importantly, EMT signalling was highly expressed among tumour epithelial but to a lesser extent in TB clusters suggests that while EMT is a general feature of the overall tumour, TB cells, seem to utilised a different set of pathways regarding their aggressive behaviour, confirming the above results.

**Figure 3.**
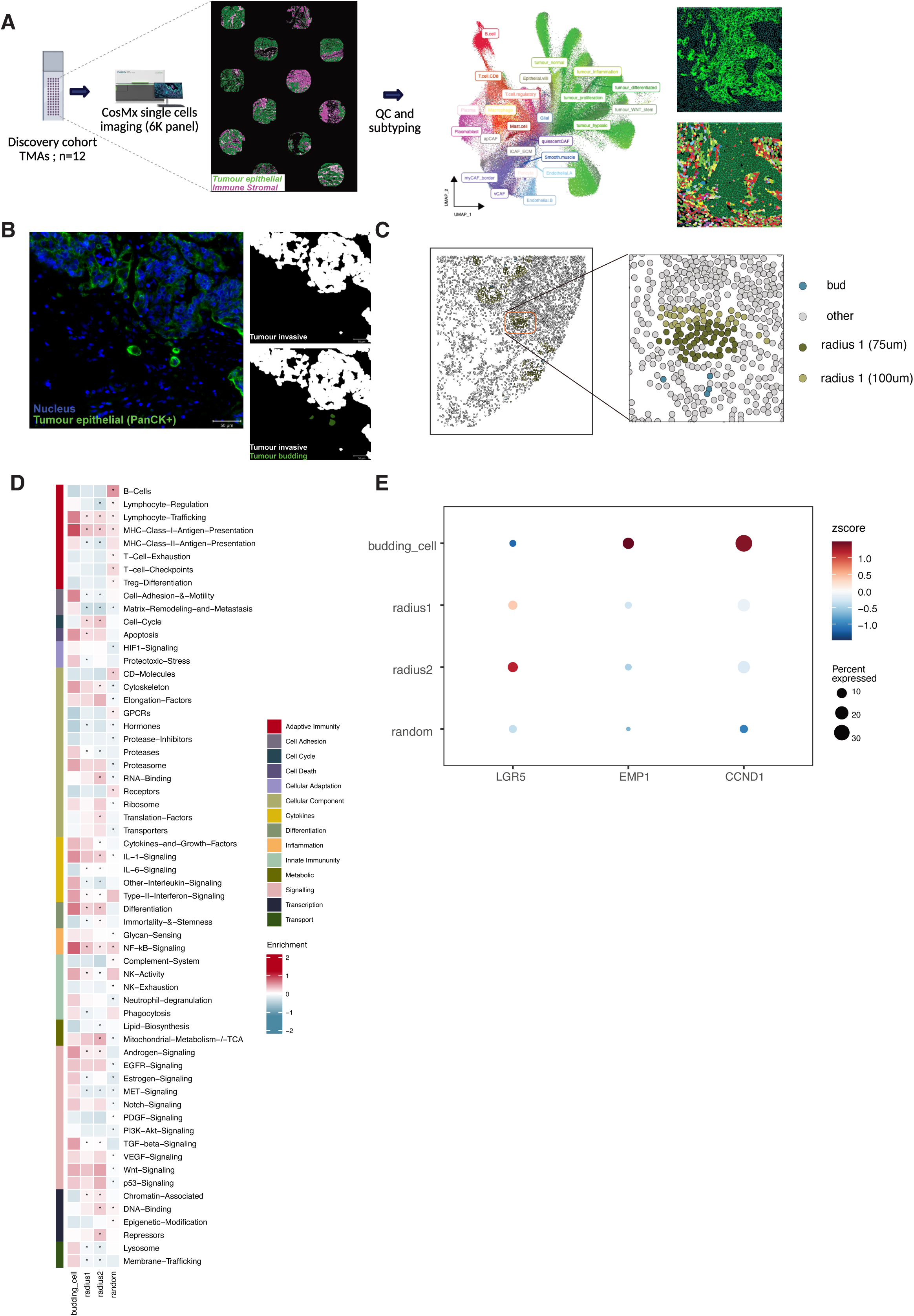
(A) The graphic illustrates CosMx single cells SMI in TMAs of discovery cohort, UMAP analysis was used to show different cell subtypes classification following quality control (QC). (B) An example of immunofluorescence staining of PanCK+ at the invasive edge area (white) and tumour budding (dark green) from single cell SMI data. (C) Adjacent invasive edge tumour was defined by radius 1 (75um), radius 2 (100um) and random cells (the other areas) to compare pathway and gene signatures to TB. (D) Pathway analysis shows the significant signalling associated with TB compared to the adjacent invasive edge tumour using the limma R package (* padj <0.05). (E) Single cell gene expression of CCND1, EMP1 and LGR5 compared between TB and adjacent invasive edge tumour cells shown in dot plot.

To thoroughly examine the molecular mechanism of budding cells and how it could be different from the adjacent tumour invasive edge (Figure 3B), radius analysis was performed (Figure 3C). Our finding demonstrates a significant increase of matrix remodeling and metastasis signature in TB cells compared to the adjacent tumour invasive edge (p<0.001). Moreover, TB cells exhibited a significant enrichment of gene signatures associated with TGF-β (p=0.001), NF-kB (p=0.001), Matrix remodelling and metastasis (p<0.001) and cell adhesion and motility (p<0.001). There was a significantly lower gene signature related to stemness (p=0.038), however, there was no significant difference in Wnt signalling, an EMT-regulating pathway, when TB cells were compared to cells in the adjacent tumour invasive edge. There was a significant increase of MHC ClassI antigen presenting (p<0.001) and IL-6 signalling (p=0.006) together with the decreased of Type-II-Interferon (p=0.001) and lymphocyte trafficking (p=0.001) indicating possible crosstalk between TB and immune cells and its ability to enhance immune evasion and diminished anti-tumour immune activity (Figure 3D).

Beyond pathway-level analysis, we examined single-gene expression within TB cells compared to their corresponding invasive edge. Based on the findings above, cyclinD1 appears to play a critical role in TB. Single cell CosMx SMI analysis revealed that TB cells expressed *CCND1* gene more than cells in the adjacent tumour invasive edge. Notably, epithelial membrane protein 1 (*EMP1*), a recognised marker of metastatic disease (27), was also elevated in TB cells, supporting their aggressive and metastatic potential. In contrast, expression of leucine-rich repeat-containing receptor 5 (*LGR5*) was reduced in TB cells (Figure 3E).

### Immune gene signatures in PanCKD regions indicate an immunosuppressive niche in tumour budding

In addition to the tumour epithelial region, regional bulk transcriptomic GeoMx DSP RNA Immune Oncology panel from the FFPE CRC full section (n=12) can also determine gene expression from non-tumour (PanCK-) cells representing the immune-microenvironment. Thus, gene expression in PanCK-invasive region was compared between low (n=6) and high (n=6) TB area (Figure 4A). When comparing low TB versus high TB phenotype in the invasive PanCK-area, *CXCL9* and *CXCL10* were significantly differentially expressed (Figure 4B-C). This suggests that expression of these chemokines may suppress TB regulation.

**Figure 4.**
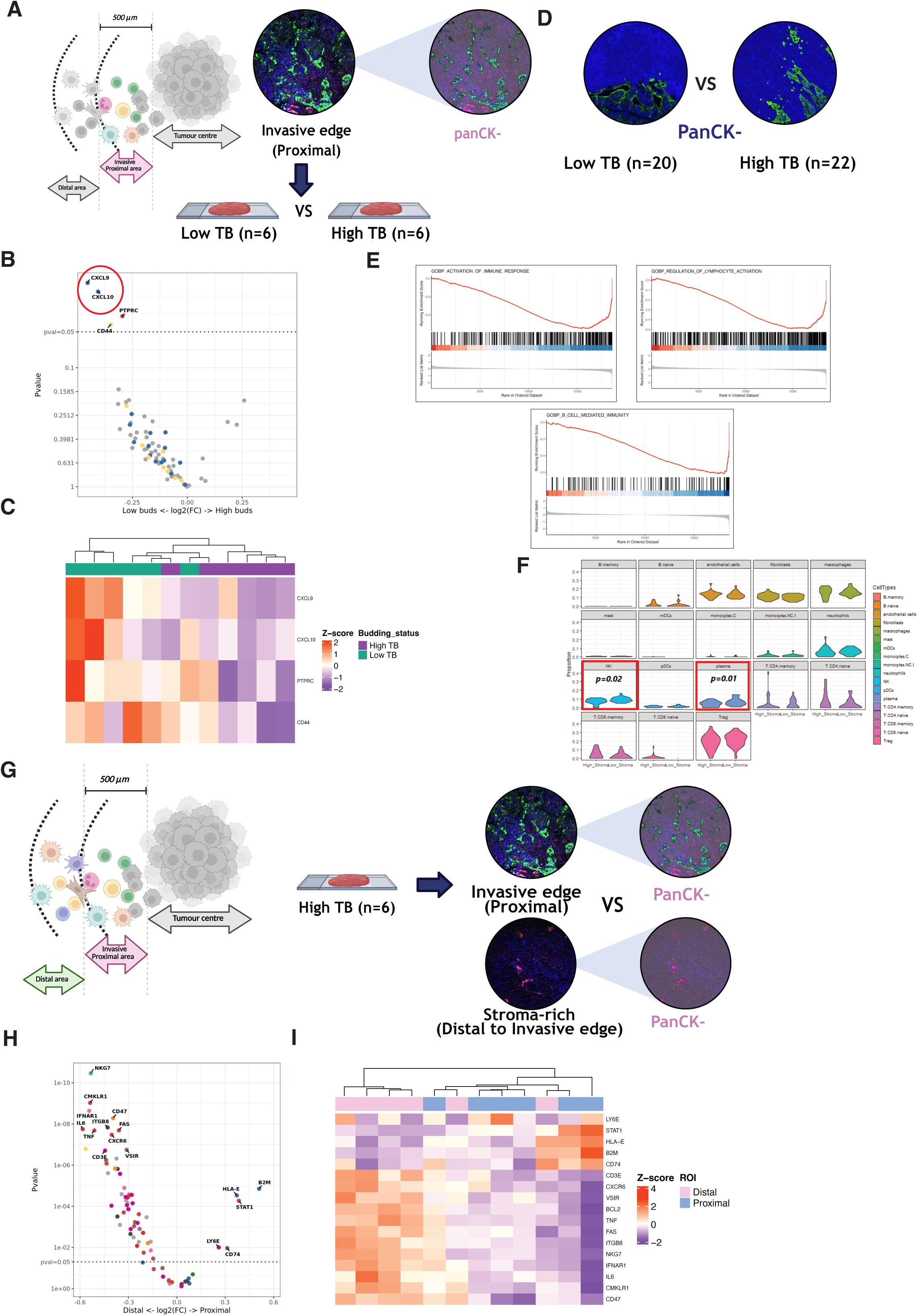
(A) Graphic shows the region of invasive non-tumour area (PanCK-) used to determine the differential genes expression compared between tumours with low (n=6) and high TB (n=6) [Note: reused figure 2A]. (B)Volcano plot revealed genes significantly expressed compared between two groups. (C) Heatmap illustrated the expression of the genes in two different groups (persian green; lowTB and purple; highTB). (D) The representative images of TMA from invasive area marked non-epithelial tissue (blue) of low (n=20) and high (n=20) TB for further analysis. (E) Enrichment plot shows the differences of genes signature associated with immune response between non-tumour area between low and high TB. (F) Cellular deconvolution for the proportion of immune cells compared between low and high TB. (G) Graphic shows the region of interested; distal and proximal stromal (PanCK-) from invasive area, used to determine the differential genes expression in tumours with high TB (n=6) [Note: reused figure 2A]. (H) Volcano plot revealed genes significantly expressed compared between two distinct areas. (I) Heatmap illustrated the expression of the genes in two different areas (pink; distal and cyan; proximal).

To further explore the findings above, gene expression from non-tumour (PanCK-) regions profiled from GeoMx DSP RNA WTA panel was also studied (Supplementary Figure S6A). Comparing invasive edge of PanCK-from low and high TB phenotypes (Figure 4D), GSEA revealed a significant decreased in gene signatures associated with immune activity in high TB invasive PanCK-area such as immune response activation (padj < 0.001, NES =-2.60), regulation of lymphocyte activation (padj <0.001, NES = –2.41) and B cell-mediated immunity (padj <0.001, NES = –2.71) compared to low TB invasive PanCK-area (Figure 4E). Gene expression deconvolution was then performed to quantify cell populations based on gene expression data. Within the PanCK-invasive area, there was a significantly lower proportion of natural killer cells (NK) (p=0.01) and plasma cells (p=0.02) (Figure 4F) in high compared to low TB cases. This may indicate that when tumours exhibited the low TB phenotype, there is an increase in inflammation whereas immunosuppressive activity is likely to be found within an invasive area of tumours with high TB.

To understand if samples with high TB are entirely devoid of immune activation or if a continuum/gradient exist, further investigation was carried out to identify gene expression between invasive proximal (closer to the budding area) and distal (distant further away from the budding area) PanCK-area using bulk transcriptomic RNA Immune Oncology GeoMx (Figure 4G). Immune-related genes, such as *NKG7, CMKLR1, IFNAR1, CD47, IL6, ITGB8, FAS, BCL2, TNF, CXCR6, CD3E, and VSIR,* were significantly expressed in the distal compared to the proximal area (Figure 4H-I).

### Spatial relationship between budding clusters and their surrounding immune cells

To further understand the role of specific immune phenotypes surrounding TB cells at the invasive margin, data from single cell CosMx SMI was analysed (Figure 5A). Immune cells were labeled after a semi-supervised method was applied (26). The nearest distance analysis was performed to identify spatial interaction between TB cells and the surrounding microenvironment (Figure 5B). Results showed that TB cells were found in close proximity to cancer associated fibroblasts (CAFs) (17.9um). Interestingly, when k-mean clustering was performed, TB clusters located either in close proximity to fibroblasts (cluster 6) or at a greater distance (cluster 22) appear to be distinct from the remainder of the budding population (Figure 5C). This observation suggests that fibroblasts—and their spatial interaction with TB cells—may play a critical role in TB development and in facilitating immune evasion and metastatic potential.

**Figure 5.**
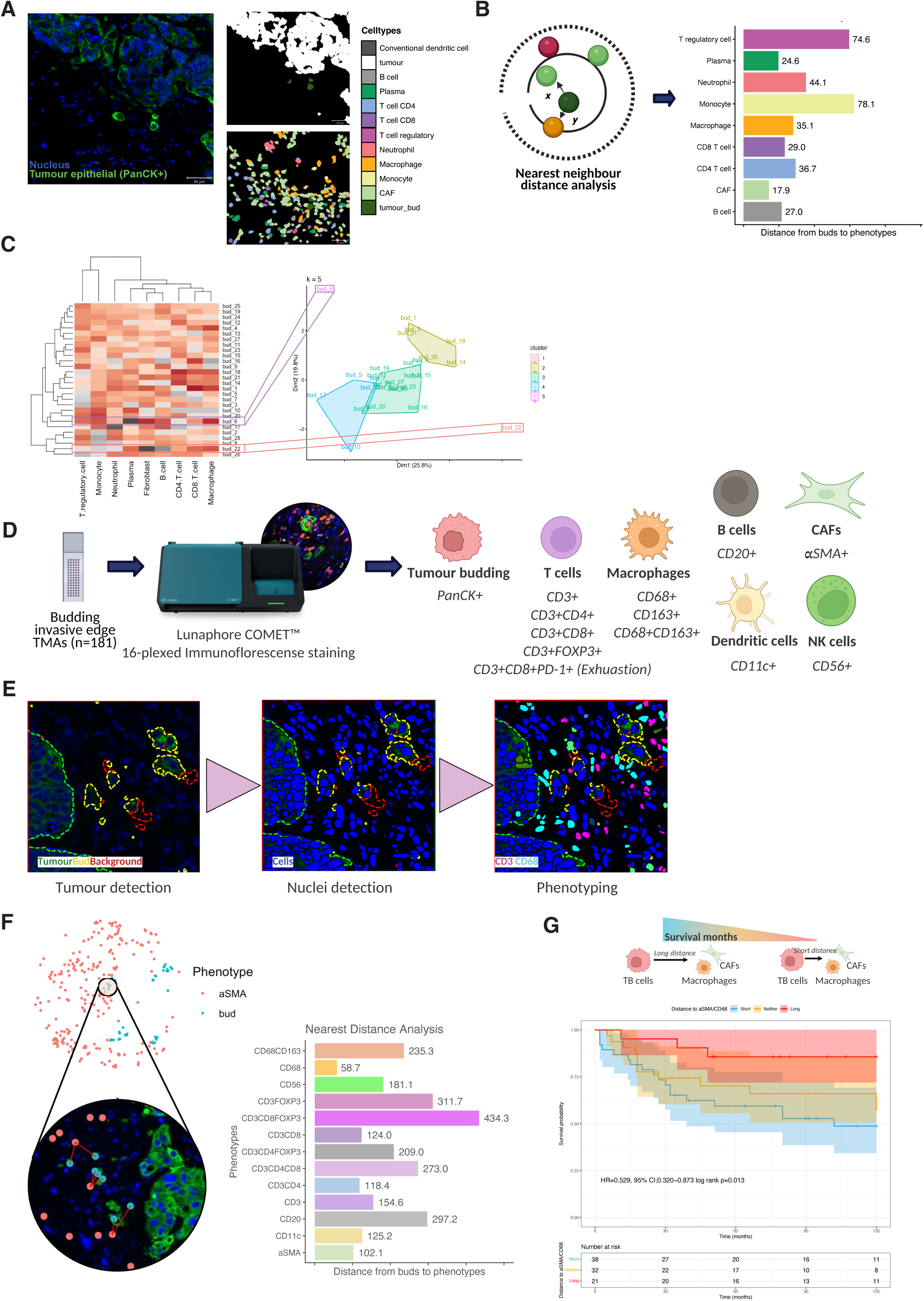
(A) An example of immunofluorescence staining of PanCK+ at the invasive edge area (white) and tumour budding (dark green) from single cell SMI data [Note: reused figure 3B]. Immune cells were subtyped using InSituType. (B) Nearest neighbour analysis shows the close proximity from TB to CAFs compared to others immune cells. (C) Heatmap illustrates the distance from TB to immune cells, importantly K-mean clustering was applied and identified distinct cluster of TB in relation to the distance to CAFs. (D) Lunaphore COMET^TM^ staining in budding invasive edge TMAs using a SPYRE panel for different immune cell phenotypes and PanCK for tumour epithelial to identify TB. (E) The workflow for image analysis; tumour detection, nuclei detection and phenotyping. (F) Nearest neighbour analysis was applied to verify the closet immune cells from TB. (G) Kaplan-Meier survival analysis based on distance from TB to aSMA+ CAFs and CD68+ macrophages for cancer specific survival (CSS) in discovery cohort.

To test this observation, a panel of selected immune markers was performed using COMET platform in budding invasive edge TMAs (n=180) (Figure 5D). After quality control of the images, TB and immune cells were phenotyped using the Visiopharm image analysis program (Figure 5E).

According to the nearest distance analysis, on average, CD68+ macrophages cells were closer to budding cells than other immune cell types, following by αSMA+ CAFs (Figure 5F). When distances were categorised as short or long relative to the TB, samples where TB were in close proximity to αSMA+ CAFs predicted significantly poorer outcomes compared to those samples with TB located farther away from αSMA+ CAFs (Supplementary Figure S11A). Interestingly, when grouped, patients with TB in close proximity to αSMA+ CAFs and CD68+ macrophages had a shorter survival compared to those TB physically distant from the both phenotypes (HR = 0.529, 95% CI: 0.320–0.873; log-rank p = 0.013) (Figure 5G).

This effect was not observed when considering distance to CD68+ cells alone, suggesting a more complex interaction between TB and its surrounding TME (Supplementary Figure S11B). Interestingly, patients with TB in close proximity to CD3+CD8+ cytotoxic T cells showed a better outcome than those with TB far from cytotoxic T cell (Supplementary Figure S12A), whereas if those T cells expressed PD-1, a marker of T cell exhaustion, patients with TB in close proximity to CD3+CD8+PD-1+ T cells exhibited worse survival compared to patients with TB far away from the phenotype (Supplementary Figure S12B).

## Discussion

Colorectal cancer (CRC) is one of the most commonly diagnosed cancers worldwide and within the UK (2). The development of screening programs has improved patient survival. However, some patients still have a poor prognosis due to metastatic disease (28). TB has recently been reported as a promising independent prognostic marker in CRC (7, 10, 29). This has resulted in an international consensus conference (ITBCC) on budding assessment, and it has been strongly recommended that TB status should be included in clinical reports in CRC (7, 30–32). A few clinical practice guidelines have included TB for CRC screening, diagnosis and treatment (33, 34). However, only a few listed TB as high-risk factor in CRC (33). In both the discovery and validation CRC patient cohorts presented in this study, high TB was demonstrated to have a strong prognostic value as well as a significant correlation with adverse clinical factors. Additionally, TB was found to be an independent prognostic factor when entering multivariate analysis in both cohorts. These results confirm that TB is a promising prognostic biomarker in CRC patients consistent with reports in other peer-reviewed studies (35).

GSEA bulk transcriptomic data and regional bulk RNA GeoMx whole transcriptomic panel revealed TNF-α and TGF-β signalling found significantly enriched in tumours with high TB. This was confirmed using single cell CosMx SMI data, when TB cells compared to the adjacent tumour invasive edge suggesting the possible regulation of these pathways within TB. The synergistic role of TNF-α and TGF-β has been used as a model to study tumour progression and metastasis (36, 37). Using laser microdissection within the invasive budding area, Li and colleagues suggested a crosstalk between the TNF-α and TGF-β pathways in the development of invasive CRC tumours (38). To further understand the molecular mechanism behind this, regional bulk RNA GeoMx immune-oncology panel was utilised, we found that *CCND1* gene, which encodes cyclinD1, was found to be highly upregulated in the invasive edge of tumours (PanCK+) with high TB. The prognostic value of cyclinD1 in CRC patients has been reported elsewhere (39). It has also been reported as a possible invasive marker in cancer, since it has been observed that cells with cyclinD1 attached to their membrane stimulate tumour invasion and metastasis in mouse model (40). There was no difference in cyclinD1 protein level when comparing the samples with low and high TB, in both tumour centre and invasive edge area. However, when assessing cyclinD1 within TB, patients expressing high cyclinD1 tend to have a poorer outcome compared to those with low cyclinD1 expression suggesting the possible role of cyclinD1 within TB leading to its aggressive behavior in CRC. Therefore, we then further investigated to confirm this finding using single cell CosMx SMI radius analysis. Results showed that TB cells expressed higher *CCND1* than cells in the adjacent tumour invasive edge, in concordance with previous studies (40, 41) suggesting cyclinD1 activity within TB may play an important role in the budding phenotype and warrants further study. Our finding demonstrates for the first time using SMI that budding cells may have different pathway activity compared to adjacent tumour invasive edge, and that cyclinD1 could be the key biomarker to target TB, this could pave the way into the development of an *in vitro* model. Moreover, we have confirmed the previous report that high *EMP1*, marker in high relapse cells (HRC), was also observed in TB (27) confirming its distinct invasive phenotype in CRC.

Studies have suggested that TB cells may not undergo proliferative activity, thus are able to avoid detection from the immune system which allows them to invade other parts of the body (42, 43). Micrometastasis cells in human-like mouse models were shown to have decreased stemness and cell proliferation (27). We have observed a similar event using SMI radius analysis as *LGR5* is lower within TB than cells in the adjacent tumour invasive edge, this could indicate that stem-like property are not acquire within the majority of TB cells as previously mentioned (44). Canellas-Socias, A., et al. reported that this phenomenon increased as the metastasis grew, suggesting a pre-metastasis stage in TB (27). Together with our finding, we hypothesised that, when produced, buds may undergo an early stage of the cell cycle, hence less proliferative and lower degree of stemness, to maintain their ability to survive undetected before extravasation.

Whether TB undergo complete or partial EMT is debatable (45, 46). Our findings show that the EMT signature was higher in PanCK+ tumours with the high TB compared to low TB phenotype at the invasive edge area. Similarly, using microdissection followed by RNA sequencing, De Smedt and colleagues reported the positive EMT gene signature in the invasive budding region compared to its main tumour centre suggesting the possibility that EMT activation regulates budding formation in CRC (47). Despite this, using known markers for mIF study, we only found a few populations of budding cells that expressed those markers suggesting that non-EMT signalling plays an important role in TB. This may also explain the partial EMT activity within TB, thus the selected markers could be crucial to understand the full activation of this pathway (45, 48, 49). Recently, using a mouse CRC model, micrometastases cells were reported to maintain expression of EPCAM and E-cadherin, suggesting both the retainment of an epithelial phenotype, and that EMT may not play an important role within these micro budding cells (27). In addition, a study by Fisher et al. suggested that the enrichment of the signature may also come from non-epithelial cells within the TME which also express genes within the EMT signature, therefore mimicking an enrichment for EMT activity (50).

In addition to pathway signalling related to TB, the analysis of regional bulk RNA GeoMx was also used to investigate gene expression within the non-tumour (PanCK-) area. Our study has reported significantly higher expression of *CXCL9* and *CXCL10* genes in low TB tumours. This is in agreement with previous reports, which observed the favorable prognostic value of these chemokines in CRC (51, 52). When a bigger sample size was studied, a significant proportion of natural killer (NK) and plasma cells were reported at the proximal invasive edge in PanCK-low compared to high TB cases. These results suggest that TB could have an immunosuppressive role which alters their microenvironment leading to disease metastasis in CRC.

To dissect the spatial crosstalk between individual TB cells and their surrounding immune phenotypes. Single cell CosMx SMI analysis, focused on TB, revealed a significant upregulation of MHC-Class I Antigen Presenting signalling within TB compared to adjacent tumour invasive edge. High expression of MHC class I molecules is well-known for its antitumorigenic properties, however, some studies reported a reverse phenotype where elevated MHC-I potentially reprograms the TME into a tumour-promoting state (53). Study from Barkal et al. showed that overexpressed MHC-I molecules can interact with NK cells and tumour-associated macrophages (TAMs) which release the “Don’t eat me” signal to inactivate the innate immune system, leading to immunosuppressive functions (54). The dynamic of immune cells activity surrounded with TB is crucial to understand if TB use the TME to evade immune surveillance.

CAFs played a crucial role in remodeling microenvironment leading to pro-metastatic (55). Recent studies show that CAFs may have different subtypes based on different gene signatures, leading to different roles in either tumour promotion or suppression (56). Our single cell CosMx SMI analysis showed that TB is found in proximity to fibroblasts, supporting the previous report that TB frequently in contact with CAFs, which leads to pro-invasion phenotype in CRC (41). Lunaphore mIF was used as an alternative method to validate these findings. There is a significant association regarding the distance from TB to αSMA+ CAFs confirming the above results that TB and CAFs may have a crucial interaction leading to the poor prognostic value of TB in CRC. Additionally, patients with TB in close proximity to both αSMA+ CAFs and CD68+ macrophages experienced the worst prognosis. A recent study also reported the engulfment of TB by CD68+ pan-macrophages indicating the crosstalk between these two cells which correlates with our observations (57). Study by Trumpi et al. suggested that co-culture patients-dervied colonosphere with macrophages induced budding formation (58). These suggested the possible role of macrophages in relation to TB development and its suppressive role in TME.

Our finding showed that T cell exhaustion (CD3+CD8+PD1+) was found proximity to TB leading to the worst outcome in CRC patients, the oppose effect showed with activated cytotoxic T cells (CD3+CD8+) suggesting the immune evasion of TB within the TME. The regulation of PD-L1/PD-1 by tumour-associated macrophages has previously been reported (59), suggesting the possible role of macrophage as a pro-tumour suppressive within TME. This mechanism is worth exploring as the finding by Lin et al. has recently reported that, within the budding area, there is an increased immune cell expression of PD-L1 leading to immune evasion of TB cells (60).

Our study has limitations. Firstly, cyclinD1 expression within TB has been assessed in high TB cases (n=180). Though single cell CosMx SMI analysis confirmed that cyclinD1 was higher in TB compared to the adjacent tumour invasive edge, however due to the small sample size, there is a need to further validate this result in a bigger cohort to confirm its prognostic value regarding the mechanism in TB cells. Additionally, as a study by Haddad et al. has recently reported pseudobudding can occur due to gland disruption and inflammation, which may misrepresent TB in CRC (42). They suggested that using PanCK staining may not represent true TB and that the pseudobudding phenotype should be taken into consideration in TB assessment. To address this, we are currently processing a larger sample set using single cell CosMx SMI for whole-transcriptome analysis, which will enable a more in-depth understanding of the true biology of TB in CRC.

In conclusion, this multi-omic investigation provides compelling evidence that TB represents a distinct epithelial phenotype with unique molecular and spatial characteristics that differ from conventional EMT-driven invasion. Our findings highlight the critical involvement of TNF-11/TGF-β signalling and cyclinD1 upregulation in TB formation, alongside the emergence of an immunosuppressive microenvironment enriched with CAFs and macrophages. These insights underscore TB as not only a robust prognostic marker but also a potential therapeutic target, offering opportunities to disrupt key signalling pathways and remodel the TME. Future studies leveraging larger cohorts and advanced spatial transcriptomics will be essential to validate these mechanisms and translate them into clinically actionable strategies aimed at improving outcomes for patients with CRC.

### Completing interests

PH performed mIF study supported by GG&C Research Endowment Fund / GN24ON047. The COMET^TM^ platform was provided as an equipment loan by Lunaphore Technologies. The authors have no other financial or personal conflicts to disclose.

## Declarations

### Ethics approval and consent to participate

For the discovery cohort, the study was approved by the West of Scotland Research Ethics Committee (REC 4: 22/ws/0207) and data are stored in the Greater Glasgow and Clyde Safehaven (SH21ON012).

### Consent for publication

Not applicable

### Availability of data and material

For the validation cohort, the data was downloaded from GEO under accession number GSE143985, published by Shinto E, Yoshida Y, Kajiwara Y, Okamoto K, Mochizuki S, Yamadera M, et al. Clinical Significance of a Gene Signature Generated from Tumor Budding Grade in Colon Cancer. DOI: 10.1245/s10434-020-08498-3 (1).

Raw data from the bulk transcriptomics data of the discovery cohort are deposited with BioProject, https://www.ncbi.nlm.nih.gov/bioproject/?term=prjna997336. Regional bulk transcriptomic and single cell spatial transcriptomic data are not publicly available due to ongoing related investigations within the research group but are available from the corresponding author upon reasonable request.

### Completing interests

PH performed mIF study supported by GG&C Research Endowment Fund / GN24ON047. The COMET^TM^ platform was provided as an equipment loan by Lunaphore Technologies. The authors have no other financial or personal conflicts to disclose.

### Authors’ contributions

PH led the project’s administration, performed the majority of the formal analysis, and wrote the main manuscript. NF conducted the formal analysis for validation cohort and assissted in drafting the manuscript. LS contributed to software development, data curation and the formal analysis. HL and AL participated in protocol development for regional bulk transcriptomic (GeoMx DSP) and AL also involved in the formal analysis of mIF. AA and AM participated in the optimisation of EMT mIF miniplex panel while JS contributed to its formal analysis. IP contributed to the methodology by providing code for mIF data analysis. CB performed the formal analysis of the validation cohort. KP and PA conducted the pathological scoring (GMS score) for the discovery cohort. CKD, and YD performed data generation for regional bulk (GeoMx DSP) and single cells (CosMx SMI) transcriptomic. JH, HM provided the tissue samples used in this study. JQ,GL,NM, AC, HW, XF, DM, OS, PD, NJ and CR provided senior infrastructure, essential resources, conceptualisation and funding acquisition. CT and JE provided high-level supervision and guide the study to publication. All authors have read and approved the final manuscript.

## List of Abbreviation

αSMA: alpha Smooth Muscle Actin
CC: Colon Cancer
CI: Confidence Interval
CRC: Colorectal Cancer
CSS: Cancer Specific Survival
CAF: Cancer Associated Fibroblast
CXCL9: C-X-C motif chemokine ligand 9
CXCL10: C-X-C motif chemokine ligand 10
DAPI: 4′,6-diamidino-2-phenylindole
DFS: Disease Free Survival
DSP: Digital Spatial Profiler
EMP: Epithelial Membrane Protein 1
EMT: Epithelial-Mesenchymal Transition
EPCAM: Epithelial Cell Adhesion Molecule
FFPE: Formalin-Fixed Paraffin-Embedded
FOV: Fields of View
GMS: Glasgow Microenvironment Score
GSEA: Gene Set Enrichment Analysis
H&E: Haematoxylin and Eosin
HR: Hazard Ratio
IHC: Immunohistochemistry
ITBCC: International Tumor Budding Consensus Conference
KM: Klintrup–Mäkinen
LGR5: Leucine-rich repeat-containing Receptor 5
mIF: Multiplex Immunofluorescence
MMR: Mismatch Repair
NES: Normalised Enrichment Score
NF-κB: Nuclear Factor kappa-light-chain-enhancer of activated B cells
PanCK: Pan-Cytokeratin
PD-1: Programme Cell Death Protein-1
ROI: Region Of Interest
SMI: Spatial Molecular Imager
TB: Tumour Budding
TNM: Tumour Node Metastasis
TMA: Tissue Microarray
TME: Tumour Microenvironment
TNF-α: Tumour Necrosis Factor alpha
TGF-β: Transforming Growth Factor beta
TSP: Tumour Stromal Percentage
UMAP: Uniform Manifold Approximation and Projection
WTA: Whole Transcriptomic Atlas

## Acknowledgements

The authors wish to acknowledge the Glasgow Tissue Research Facility and NHS Greater Glasgow and Clyde biorepository for helping in preparing the tissue samples for this study. The authors would also like to acknowledge Dr. Suyanee Thongchot for her accommodation and the Siriraj Research Fund, Faculty of Medicine Siriraj Hospital for the financial support (R016533013). The authors would also like to acknowledge the Academic unit of Surgery at the Glasgow Royal Infirmary for access to human cohort. The authors wish to acknowledge Dr. Catherine Winchester for help reviewing the manuscript prior to submission.

## Authors’ information (optional)

Not applicable

## References

1. Shinto E, Yoshida Y, Kajiwara Y, Okamoto K, Mochizuki S, Yamadera M, et al. Clinical Significance of a Gene Signature Generated from Tumor Budding Grade in Colon Cancer. Ann Surg Oncol. 2020;27(10):4044–54.

2. Rawla P, Sunkara T, Barsouk A. Epidemiology of colorectal cancer: incidence, mortality, survival, and risk factors. Prz Gastroenterol. 2019;14(2):89–103.

3. Navarro M, Nicolas A, Ferrandez A, Lanas A. Colorectal cancer population screening programs worldwide in 2016: An update. World J Gastroenterol. 2017;23(20):3632–42.

4. Vayrynen V, Wirta EV, Seppala T, Sihvo E, Mecklin JP, Vasala K, et al. Incidence and management of patients with colorectal cancer and synchronous and metachronous colorectal metastases: a population-based study. BJS Open. 2020;4(4):685–92.

5. Biller LH, Schrag D. Diagnosis and Treatment of Metastatic Colorectal Cancer: A Review. JAMA. 2021;325(7):669–85.

6. Xie YH, Chen YX, Fang JY. Comprehensive review of targeted therapy for colorectal cancer. Signal Transduct Target Ther. 2020;5(1):22.

7. Lugli A, Kirsch R, Ajioka Y, Bosman F, Cathomas G, Dawson H, et al. Recommendations for reporting tumor budding in colorectal cancer based on the International Tumor Budding Consensus Conference (ITBCC) 2016. Mod Pathol. 2017;30(9):1299–311.

8. Zlobec I, Hädrich M, Dawson H, Koelzer VH, Borner M, Mallaev M, et al. Intratumoural budding (ITB) in preoperative biopsies predicts the presence of lymph node and distant metastases in colon and rectal cancer patients. British Journal of Cancer. 2013;110(4):1008–13.

9. Koelzer VH, Zlobec I, Lugli A. Tumor budding in colorectal cancer--ready for diagnostic practice? Hum Pathol. 2016;47(1):4–19.

10. Lugli A, Zlobec I, Berger MD, Kirsch R, Nagtegaal ID. Tumour budding in solid cancers. Nat Rev Clin Oncol. 2021;18(2):101–15.

11. Salhia B, Trippel M, Pfaltz K, Cihoric N, Grogg A, Ladrach C, et al. High tumor budding stratifies breast cancer with metastatic properties. Breast Cancer Res Treat. 2015;150(2):363–71.

12. Pai RK, Cheng YW, Jakubowski MA, Shadrach BL, Plesec TP, Pai RK. Colorectal carcinomas with submucosal invasion (pT1): analysis of histopathological and molecular factors predicting lymph node metastasis. Mod Pathol. 2017;30(1):113–22.

13. Slik K, Blom S, Turkki R, Välimäki K, Kurki S, Mustonen H, et al. Combined epithelial marker analysis of tumour budding in stage II colorectal cancer. The Journal of Pathology: Clinical Research. 2019;5(1):63–78.

14. Ueno H, Ishiguro M, Nakatani E, Ishikawa T, Uetake H, Matsuda C, et al. Prospective Multicenter Study on the Prognostic and Predictive Impact of Tumor Budding in Stage II Colon Cancer: Results From the SACURA Trial. J Clin Oncol. 2019;37(22):1886–94.

15. Mitrovic B, Handley K, Assarzadegan N, Chang HL, Dawson HAE, Grin A, et al. Prognostic and Predictive Value of Tumor Budding in Colorectal Cancer. Clin Colorectal Cancer. 2021;20(3):256–64.

16. Hatthakarnkul P, Quinn JA, Ammar A, Lynch G, Van Wyk H, McMillan DC, et al. Molecular mechanisms of tumour budding and its association with microenvironment in colorectal cancer. Clin Sci (Lond). 2022;136(8):521–35.

17. Hatthakarnkul P, Quinn JA, Matly AAM, Ammar A, van Wyk HC, McMillan DC, et al. Systematic review of tumour budding and association with common mutations in patients with colorectal cancer. Crit Rev Oncol Hematol. 2021;167:103490.

18. McMillan DC. The systemic inflammation-based Glasgow Prognostic Score: a decade of experience in patients with cancer. Cancer Treat Rev. 2013;39(5):534–40.

19. Lu X, Guo W, Xu W, Zhang X, Shi Z, Zheng L, et al. Prognostic value of the Glasgow prognostic score in colorectal cancer: a meta-analysis of 9,839 patients. Cancer Manag Res. 2019;11:229–49.

20. Proctor MJ, Morrison DS, Talwar D, Balmer SM, O’Reilly DS, Foulis AK, et al. An inflammation-based prognostic score (mGPS) predicts cancer survival independent of tumour site: a Glasgow Inflammation Outcome Study. Br J Cancer. 2011;104(4):726–34.

21. van Wyk HC, Park JH, Edwards J, Horgan PG, McMillan DC, Going JJ. The relationship between tumour budding, the tumour microenvironment and survival in patients with primary operable colorectal cancer. Br J Cancer. 2016;115(2):156–63.

22. Yeakley JM, Shepard PJ, Goyena DE, VanSteenhouse HC, McComb JD, Seligmann BE. A trichostatin A expression signature identified by TempO-Seq targeted whole transcriptome profiling. PloS one. 2017;12(5):e0178302.

23. Dobin A, Davis CA, Schlesinger F, Drenkow J, Zaleski C, Jha S, et al. STAR: ultrafast universal RNA-seq aligner. Bioinformatics. 2013;29(1):15–21.

24. Alexander PG, Roseweir AK, Pennel KAF, van Wyk HC, Powell A, McMillan DC, et al. The Glasgow Microenvironment Score associates with prognosis and adjuvant chemotherapy response in colorectal cancer. Br J Cancer. 2021;124(4):786–96.

25. Matthews L, Gopinath G, Gillespie M, Caudy M, Croft D, de Bono B, et al. Reactome knowledgebase of human biological pathways and processes. Nucleic Acids Res. 2009;37(Database issue):D619–22.

26. Danaher P, Zhao E, Yang Z, Ross D, Gregory M, Reitz Z, et al. Insitutype: likelihood-based cell typing for single cell spatial transcriptomics. bioRxiv preprint. 2022.

27. Canellas-Socias A, Cortina C, Hernando-Momblona X, Palomo-Ponce S, Mulholland EJ, Turon G, et al. Metastatic recurrence in colorectal cancer arises from residual EMP1(+) cells. Nature. 2022;611(7936):603–13.

28. Zeineddine FA, Zeineddine MA, Yousef A, Gu Y, Chowdhury S, Dasari A, et al. Survival improvement for patients with metastatic colorectal cancer over twenty years. NPJ Precis Oncol. 2023;7(1):16.

29. Sirin AH, Sokmen S, Unlu SM, Ellidokuz H, Sarioglu S. The prognostic value of tumor budding in patients who had surgery for rectal cancer with and without neoadjuvant therapy. Tech Coloproctol. 2019;23(4):333–42.

30. Haddad TS, Lugli A, Aherne S, Barresi V, Terris B, Bokhorst JM, et al. Improving tumor budding reporting in colorectal cancer: a Delphi consensus study. Virchows Arch. 2021;479(3):459–69.

31. Graham RP, Vierkant RA, Tillmans LS, Wang AH, Laird PW, Weisenberger DJ, et al. Tumor Budding in Colorectal Carcinoma: Confirmation of Prognostic Significance and Histologic Cutoff in a Population-based Cohort. Am J Surg Pathol. 2015;39(10):1340–6.

32. Zlobec I, Berger MD, Lugli A. Tumour budding and its clinical implications in gastrointestinal cancers. Br J Cancer. 2020;123(5):700–8.

33. Argiles G, Tabernero J, Labianca R, Hochhauser D, Salazar R, Iveson T, et al. Localised colon cancer: ESMO Clinical Practice Guidelines for diagnosis, treatment and follow-up. Ann Oncol. 2020;31(10):1291–305.

34. Watanabe T, Muro K, Ajioka Y, Hashiguchi Y, Ito Y, Saito Y, et al. Japanese Society for Cancer of the Colon and Rectum (JSCCR) guidelines 2016 for the treatment of colorectal cancer. Int J Clin Oncol. 2018;23(1):1–34.

35. Rogers AC, Winter DC, Heeney A, Gibbons D, Lugli A, Puppa G, et al. Systematic review and meta-analysis of the impact of tumour budding in colorectal cancer. Br J Cancer. 2016;115(7):831–40.

36. Liu ZW, Zhang YM, Zhang LY, Zhou T, Li YY, Zhou GC, et al. Duality of Interactions Between TGF-beta and TNF-alpha During Tumor Formation. Front Immunol. 2021;12:810286.

37. Hahn S, Nam MO, Noh JH, Lee DH, Han HW, Kim DH, et al. Organoid-based epithelial to mesenchymal transition (OEMT) model: from an intestinal fibrosis perspective. Sci Rep. 2017;7(1):2435.

38. Li H, Zhong A, Li S, Meng X, Wang X, Xu F, et al. The integrated pathway of TGFbeta/Snail with TNFalpha/NFkappaB may facilitate the tumor-stroma interaction in the EMT process and colorectal cancer prognosis. Sci Rep. 2017;7(1):4915.

39. Li Y, Wei J, Xu C, Zhao Z, You T. Prognostic significance of cyclin D1 expression in colorectal cancer: a meta-analysis of observational studies. PLoS One. 2014;9(4):e94508.

40. Fuste NP, Castelblanco E, Felip I, Santacana M, Fernandez-Hernandez R, Gatius S, et al. Characterization of cytoplasmic cyclin D1 as a marker of invasiveness in cancer. Oncotarget. 2016;7(19):26979–91.

41. Garcia-Rodriguez JL, Korsgaard U, Vissing SM, Paasch TP, Semenova M, Vendelbo SL, et al. Cancer-associated fibroblasts shape the formation of budding cancer cells at the invasive front of human colorectal cancer. Commun Biol. 2025;8(1):1345.

42. Haddad TS, van den Dobbelsteen L, Ozturk SK, Geene R, Nijman IJ, Verrijp K, et al. Pseudobudding: ruptured glands do not represent true tumor buds. J Pathol. 2023;261(1):19–27.

43. Dawson H, Koelzer VH, Karamitopoulou E, Economou M, Hammer C, Muller DE, et al. The apoptotic and proliferation rate of tumour budding cells in colorectal cancer outlines a heterogeneous population of cells with various impacts on clinical outcome. Histopathology. 2014;64(4):577–84.

44. Sato K, Uehara T, Iwaya M, Nakajima T, Miyagawa Y, Ota H, et al. Correlation of clinicopathological features and LGR5 expression in colon adenocarcinoma. Ann Diagn Pathol. 2020;48:151587.

45. Grigore AD, Jolly MK, Jia D, Farach-Carson MC, Levine H. Tumor Budding: The Name is EMT. Partial EMT. J Clin Med. 2016;5(5).

46. Pavlic A, Bostjancic E, Kavalar R, Ilijevec B, Bonin S, Zanconati F, et al. Tumour budding and poorly differentiated clusters in colon cancer – different manifestations of partial epithelial-mesenchymal transition. J Pathol. 2022;258(3):278–88.

47. De Smedt L, Palmans S, Andel D, Govaere O, Boeckx B, Smeets D, et al. Expression profiling of budding cells in colorectal cancer reveals an EMT-like phenotype and molecular subtype switching. Br J Cancer. 2017;116(1):58–65.

48. Yang J, Antin P, Berx G, Blanpain C, Brabletz T, Bronner M, et al. Guidelines and definitions for research on epithelial-mesenchymal transition. Nat Rev Mol Cell Biol. 2020;21(6):341–52.

49. Simeonov KP, Byrns CN, Clark ML, Norgard RJ, Martin B, Stanger BZ, et al. Single-cell lineage tracing of metastatic cancer reveals selection of hybrid EMT states. Cancer Cell. 2021;39(8):1150–62 e9.

50. Fisher NC, Byrne RM, Leslie H, Wood C, Legrini A, Cameron AJ, et al. Biological Misinterpretation of Transcriptional Signatures in Tumor Samples Can Unknowingly Undermine Mechanistic Understanding and Faithful Alignment with Preclinical Data. Clin Cancer Res. 2022;28(18):4056–69.

51. Chen J, Ye X, Pitmon E, Lu M, Wan J, Jellison ER, et al. IL-17 inhibits CXCL9/10-mediated recruitment of CD8(+) cytotoxic T cells and regulatory T cells to colorectal tumors. J Immunother Cancer. 2019;7(1):324.

52. Zhang J, Tao J, Gao RN, Wei ZY, He YS, Ren CY, et al. Cytotoxic T-Cell Trafficking Chemokine Profiles Correlate With Defined Mucosal Microbial Communities in Colorectal Cancer. Front Immunol. 2021;12:715559.

53. Wu X, Li T, Jiang R, Yang X, Guo H, Yang R. Targeting MHC-I molecules for cancer: function, mechanism, and therapeutic prospects. Mol Cancer. 2023;22(1):194.

54. Barkal AA, Weiskopf K, Kao KS, Gordon SR, Rosental B, Yiu YY, et al. Engagement of MHC class I by the inhibitory receptor LILRB1 suppresses macrophages and is a target of cancer immunotherapy. Nat Immunol. 2018;19(1):76–84.

55. Lan X, Li W, Zhao K, Wang J, Li S, Zhao H. Revisiting the role of cancer-associated fibroblasts in tumor microenvironment. Front Immunol. 2025;16:1582532.

56. Cords L, de Souza N, Bodenmiller B. Classifying cancer-associated fibroblasts-The good, the bad, and the target. Cancer Cell. 2024;42(9):1480–5.

57. Koelzer VH, Canonica K, Dawson H, Sokol L, Karamitopoulou-Diamantis E, Lugli A, et al. Phenotyping of tumor-associated macrophages in colorectal cancer: Impact on single cell invasion (tumor budding) and clinicopathological outcome. Oncoimmunology. 2016;5(4):e1106677.

58. Trumpi K, Frenkel N, Peters T, Korthagen NM, Jongen JMJ, Raats D, et al. Macrophages induce “budding” in aggressive human colon cancer subtypes by protease-mediated disruption of tight junctions. Oncotarget. 2018;9(28):19490–507.

59. Pu Y, Ji Q. Tumor-Associated Macrophages Regulate PD-1/PD-L1 Immunosuppression. Front Immunol. 2022;13:874589.

60. Lin JR, Wang S, Coy S, Chen YA, Yapp C, Tyler M, et al. Multiplexed 3D atlas of state transitions and immune interaction in colorectal cancer. Cell. 2023;186(2):363–81 e19.

