## Supplementary figure legend for "Epithelial Plasticity and an Immune Suppressive Microenvironment Underpin Tumour Budding in Colorectal Cancer"

***Supplementary Figure S1*** *Tumour cells clusters identified using InsituType across FOVs*

***Supplementary Figure S2*** *(A-I) Kaplan-Meier survival analysis based on tumour budding phenotype stratified by clinical factors for cancer specific survival (CSS) in discovery cohort.*

***Supplementary Figure S3*** *(A) Kaplan-Meier survival analysis based on TB status for disease-free survival (DFS) in CRC patients of the discovery cohort. Kaplan-Meier survival analysis based on TB status stratified by disease’s type; Colon, Rectal, for (B) cancer-specific survival (CSS) and (C) disease-free survival (DFS) in CRC patients of the discovery cohort.*

**Supplementary Figure S4** (A) Kaplan-Meier survival analysis based on tumour budding phenotype stratified by clinical factors for disease-free survival (DFS) in validated CRC cohort (n=85.). (B) The correlation plot showed the standardised residual of data; positive associations are in blue and no association in orange, the bigger size of the circle the more significant association was found. (C) Kaplan-Meier survival analysis based on tumour budding phenotype stratified by clinical factors for disease-free survival (DFS) in validation cohort.

***Supplementary Figure S5*** *(A) Simplified graphic summarising the number of patients in discovery and validation cohort used in transcriptomic analysis. Dot plot based GSEA analysis shows top 10 gene sets significantly up and down-regulated in (B, G) hallmark and (D, I) curated gene set pathways. (C,E,H and J) Enrichment plot shows the differences of signalling pathways involved between tumours with low and high TB in both discovery and validation cohorts.*

***Supplementary Figure S6*** *(A) Graphic summarising the number of patients in discovery cohort used in regional bulk RNA GeoMx (WTA panel). (B) The representative images of TMA from invasive area marked epithelial area (PanCK+; red) with low (n=20) and high (n=23) TB for further analysis. (C) Dot plot based GSEA analysis shows top 10 gene sets significantly up and down-regulated in hallmark gene set pathways. (D) Enrichment plot shows the differences of signalling pathways involved between tumours with low and high TB. (E) Multiplex staining showed the expression of an E-cadherin and β-catenin within TB cells (F) The percentage of budding population that expressed high E-cadherin- β-catenin+*

***Supplementary Figure S7*** *Boxplot shows continuous weighted histoscore of cyclinD1 protein expression compared between tumour with high and low TB assessed within (A) tumour centre and (B) invasive edge areas.t-test applied for comparing between two groups.*

***Supplementary Figure S8*** *The simplified diagram describes the pre-processing steps of single cell CosMx SMI data*

***Supplementary Figure S9*** *The bar plot shows the gene signature expressed within TB, which represents a different type of tumour signature using InSituType*

*.*

***Supplementary Figure S10*** *The heatmap illustrates the comparison of pathway activity in different tumour type compared to TB*

***Supplementary Figure S11*** *(A) Kaplan-Meier survival analysis based on the distance from TB to aSMA+ CAFs for cancer specific survival (CSS) in discovery cohort. (B) Kaplan-Meier survival analysis based on the distance from TB to CD68+ macrophages for cancer specific survival (CSS) in discovery cohort.*

***Supplementary Figure S12*** *(A) Kaplan-Meier survival analysis based on the distance from TB to CD3+CD8+ cytotoxic T cells for cancer specific survival (CSS) in discovery cohort. (B) Kaplan-Meier survival analysis based on the distance from TB to CD3+CD8+PD1+ T cell exhaustion for cancer specific survival (CSS) in discovery cohort.*
