## Supplementary figures and images for "Epithelial Plasticity and an Immune Suppressive Microenvironment Underpin Tumour Budding in Colorectal Cancer"

### supplementary figure1

***Supplementary Figure 1***

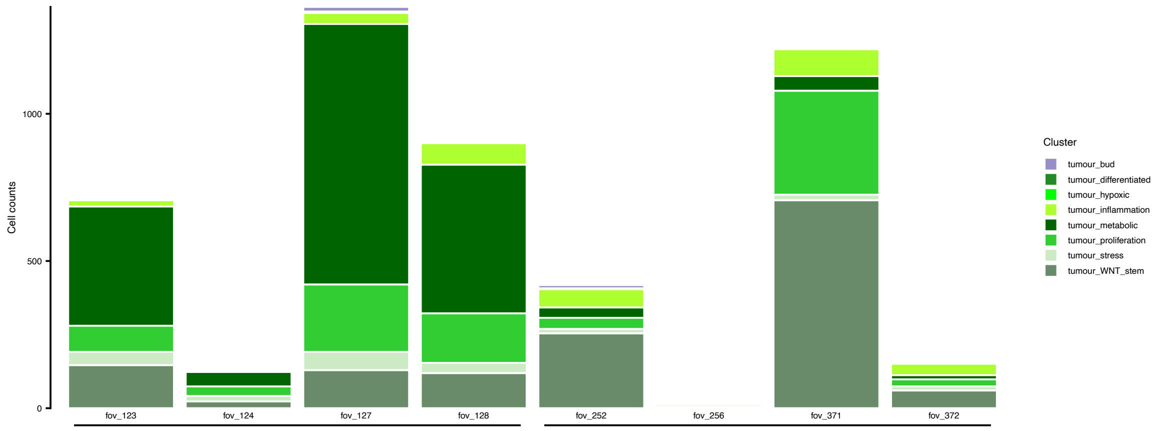

### supplementary figure2

*Supplementary Figure 2*

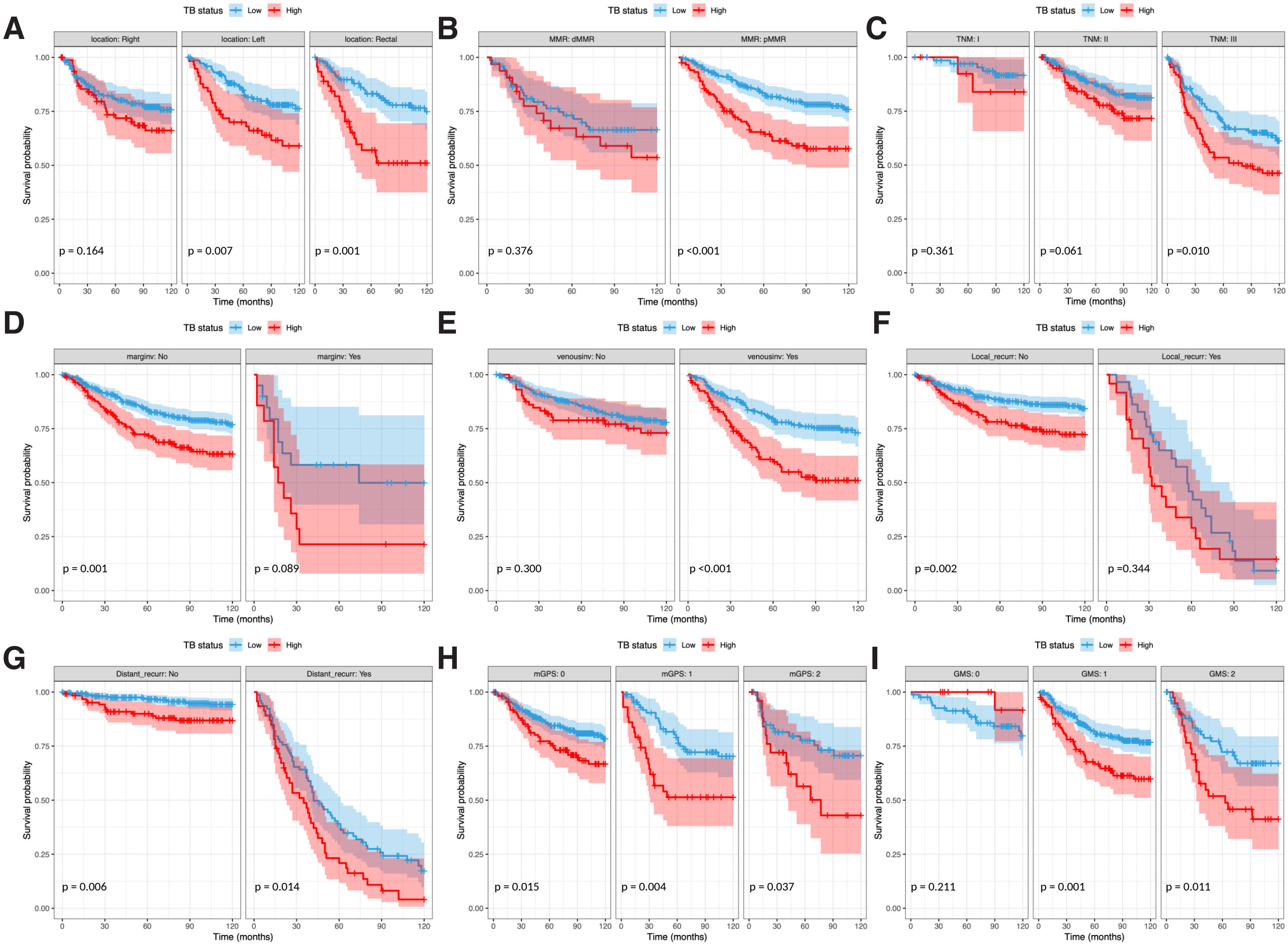

### supplementary figure3

##### *Supplementary Figure 3*

A

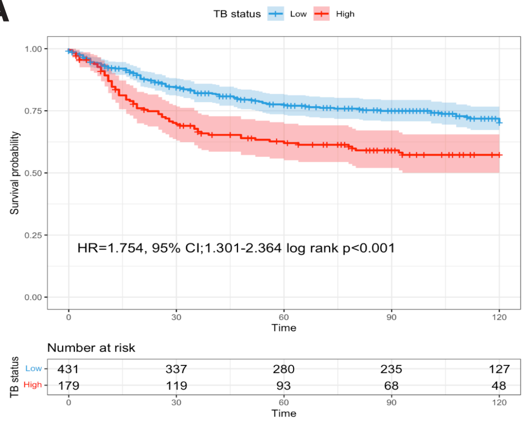


B

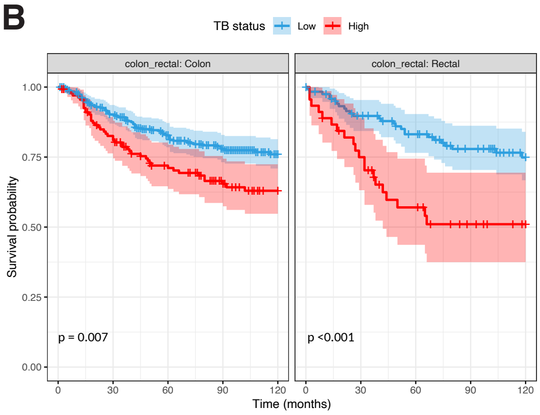


C

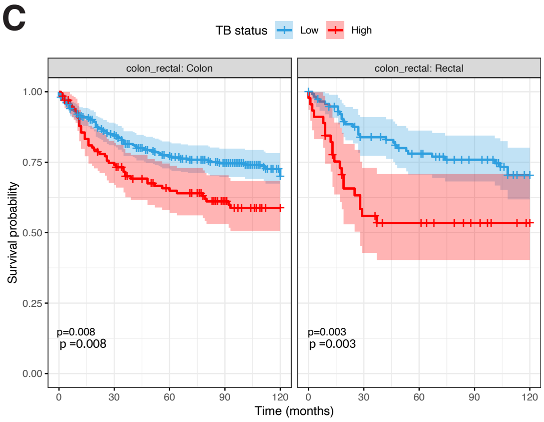

### supplementary figure4

*Supplementary Figure4*

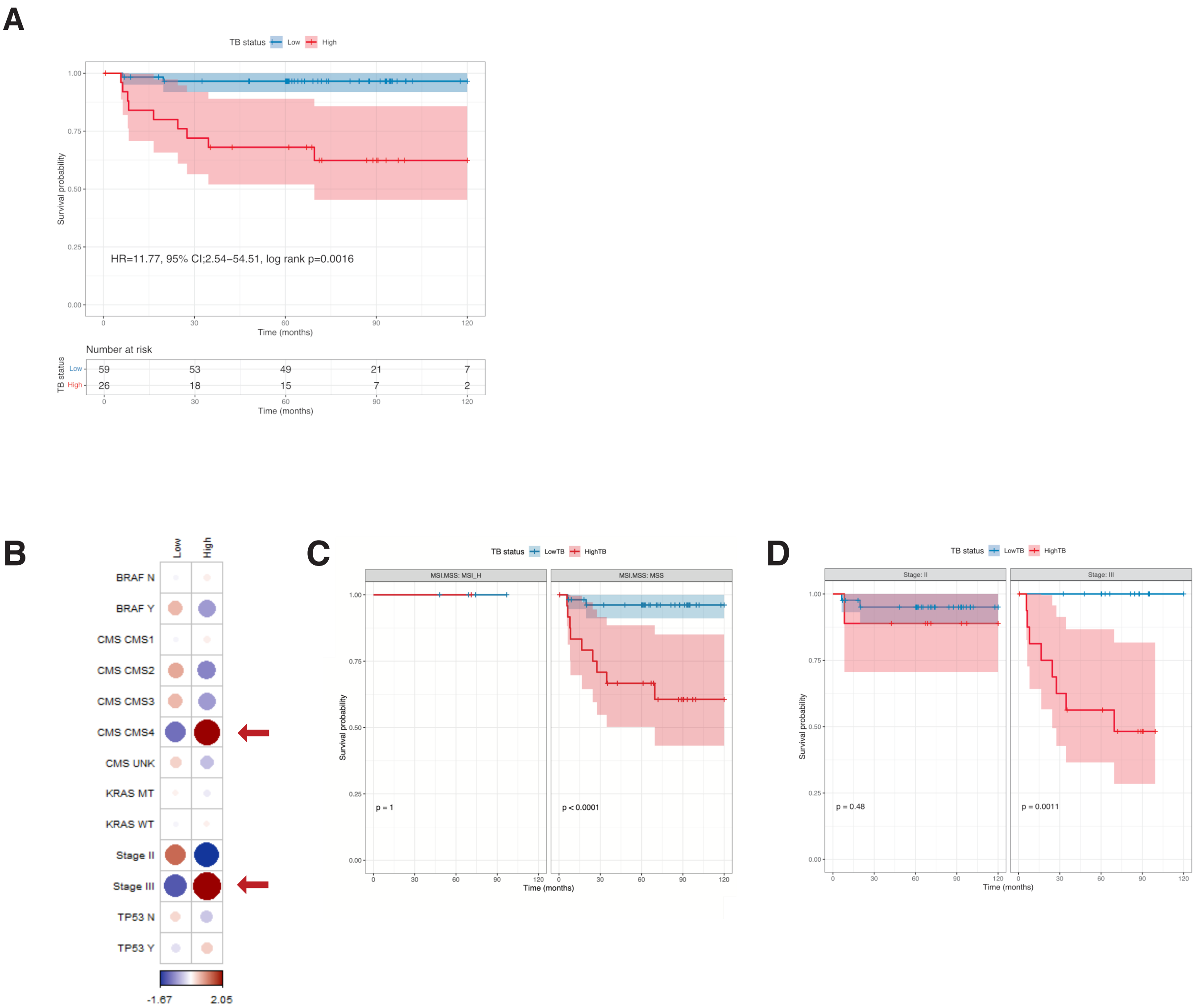

### supplementary figure5

*Supplementary Figure 5*

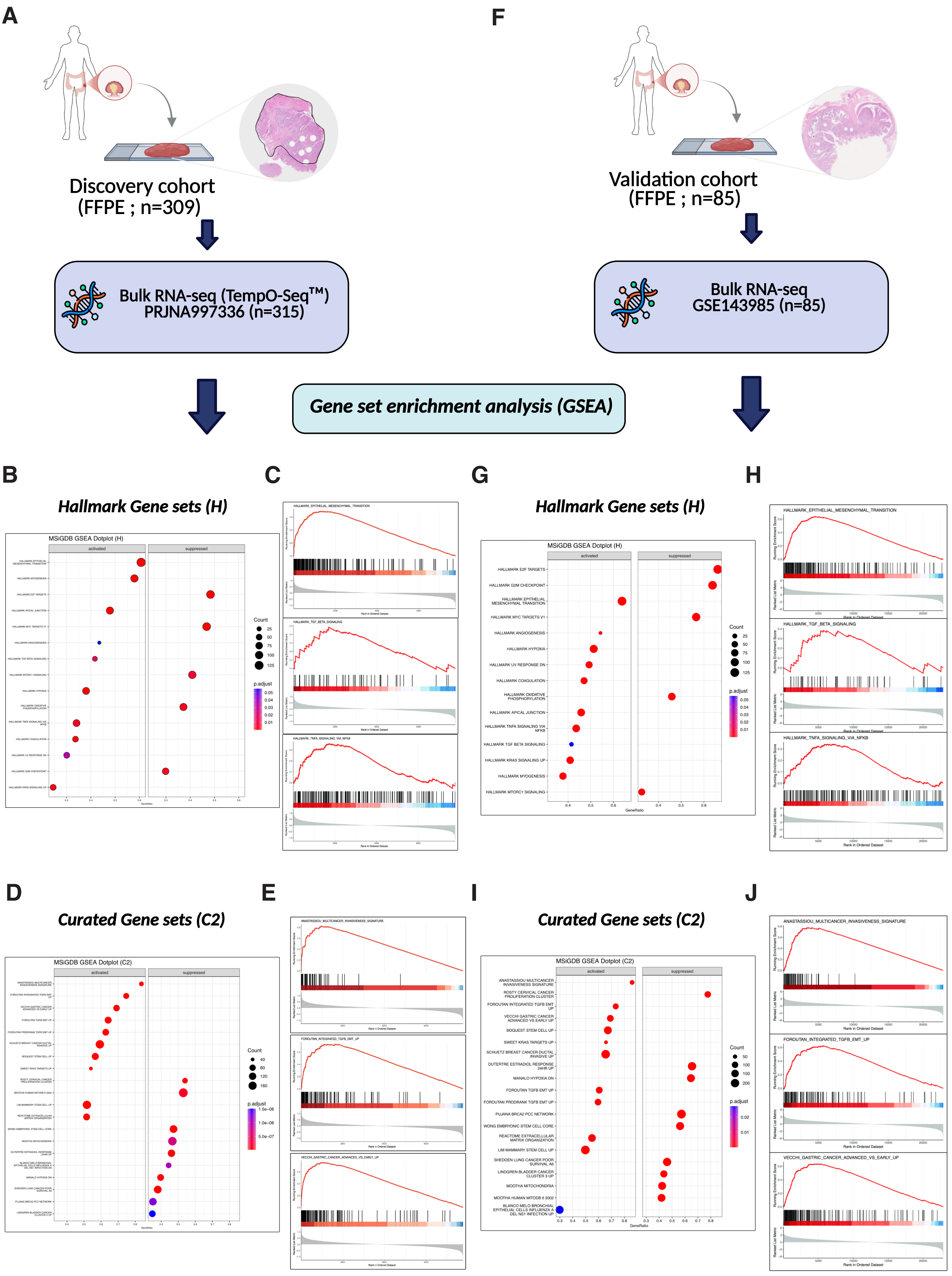

### supplementary figure6

##### *Supplementary Figure 6*

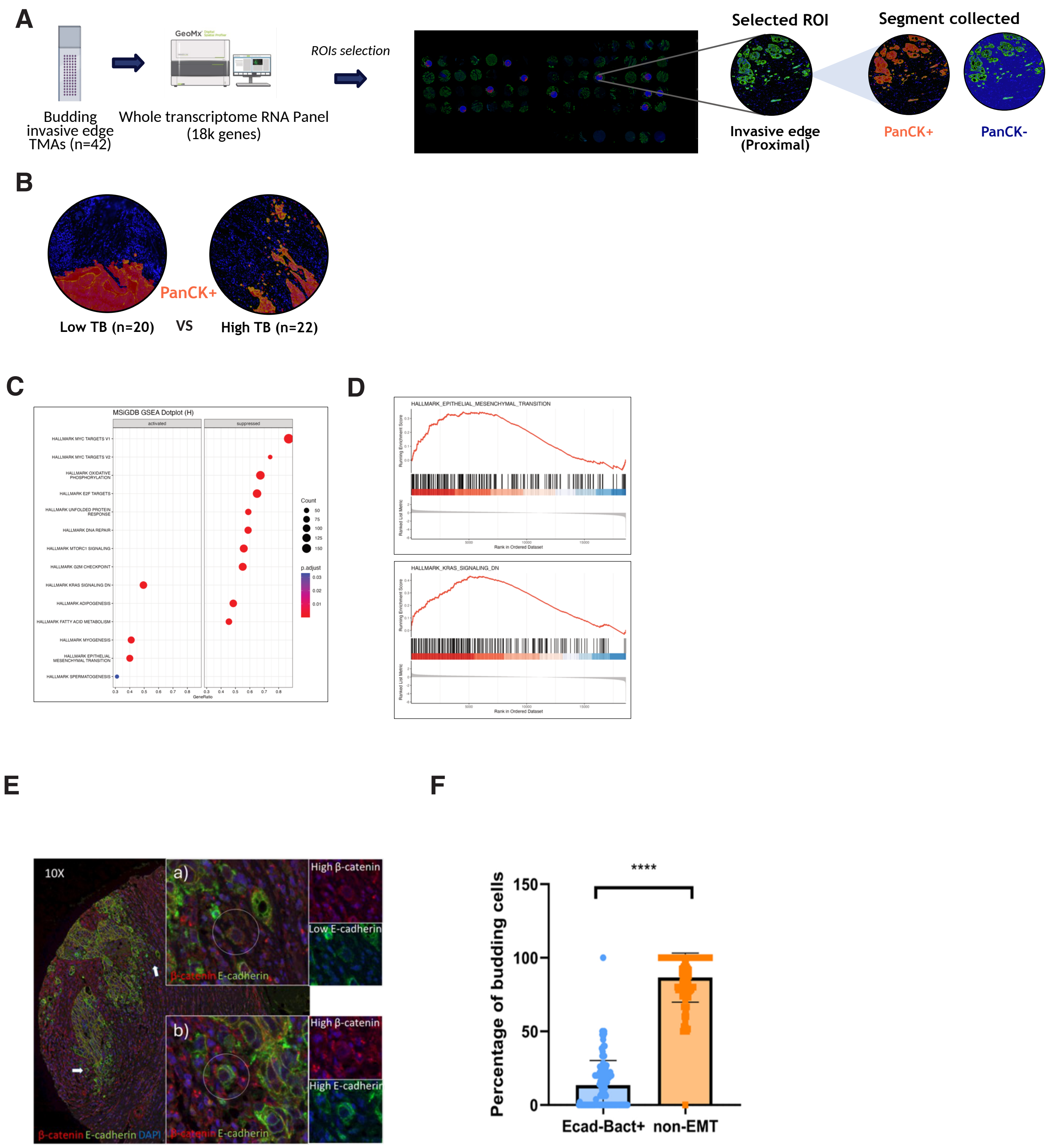

### supplementary figure7

*Supplementary Figure 7*

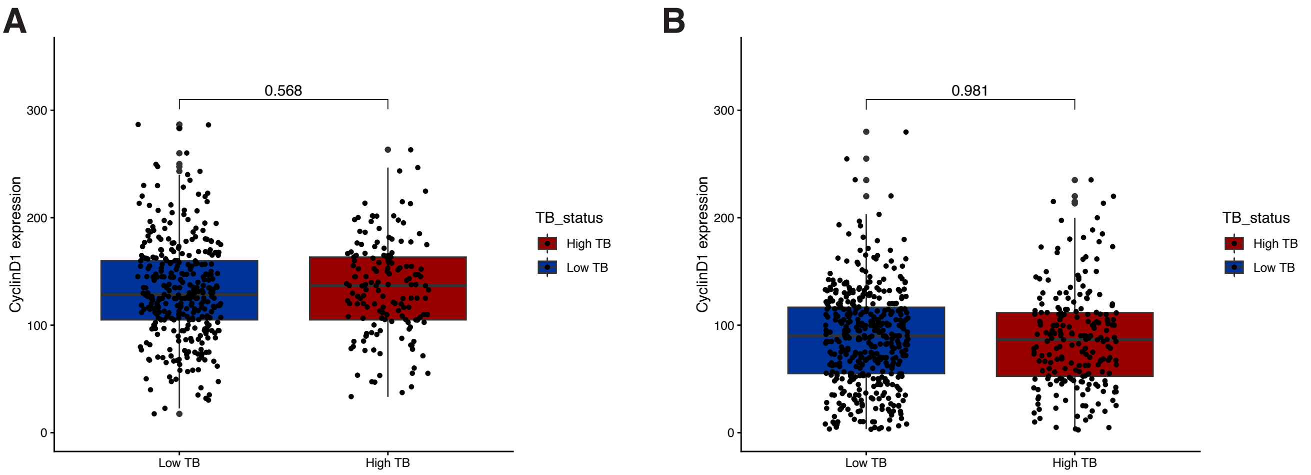

### supplementary figure9

#### *Supplementary Figure 9*

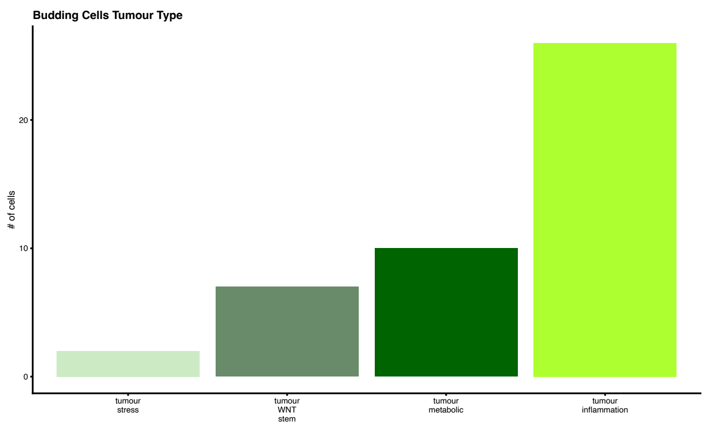

### supplementary figure10

*Supplementary Figure 10*

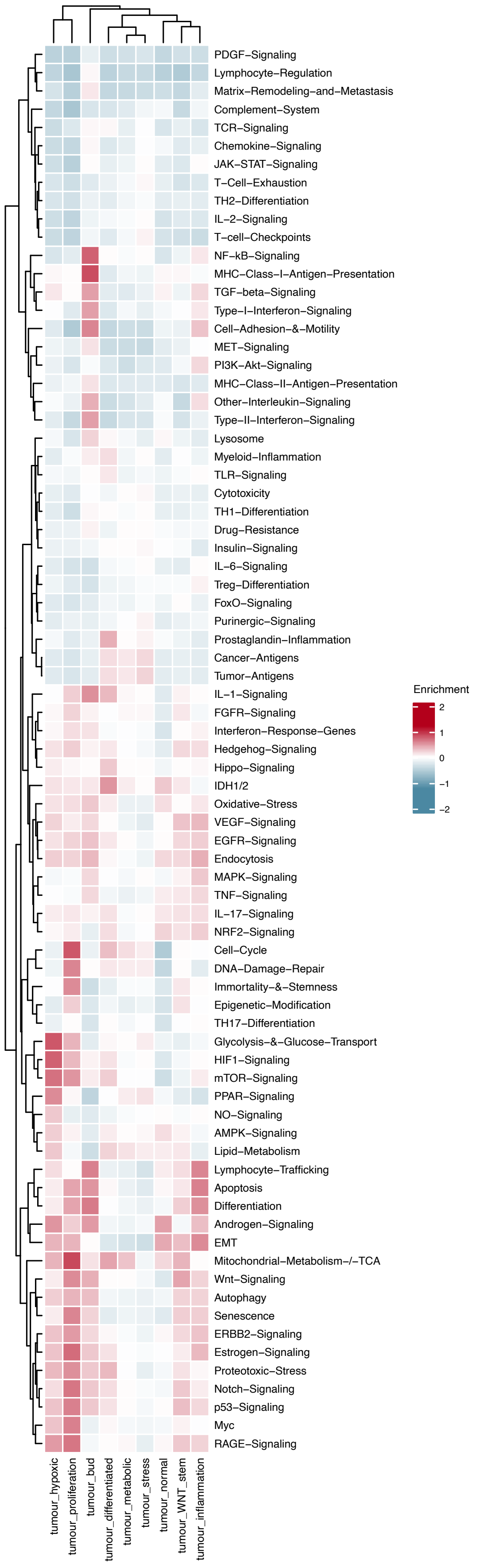

### supplementary figure11

##### *Supplementary Figure 11*

A

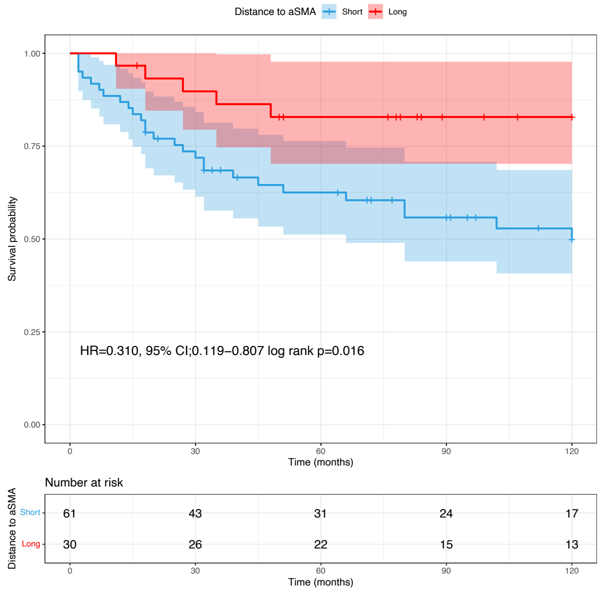


B

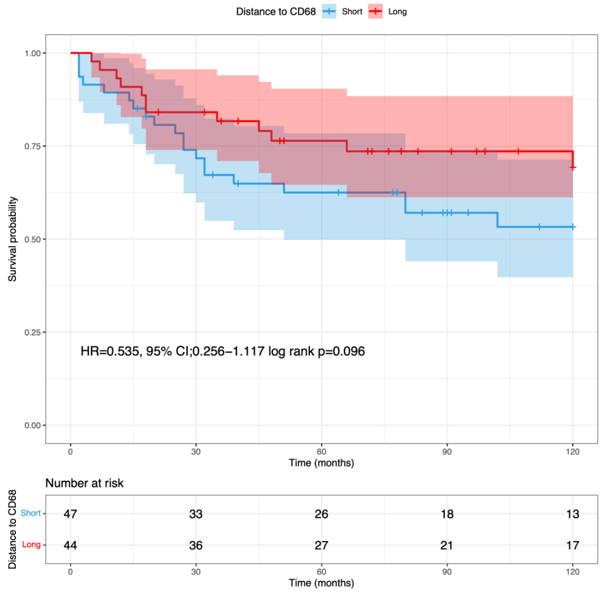

### supplementary figure12

#### Supplementary Figure 12

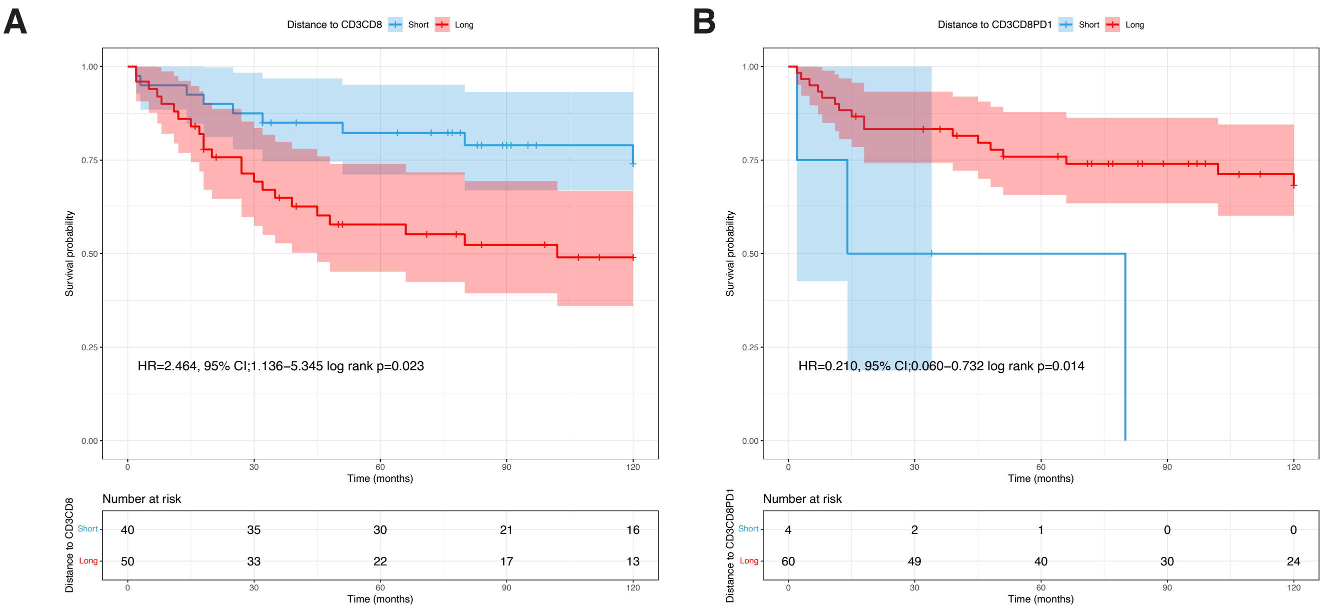
