## supplementary figure8 for "Epithelial Plasticity and an Immune Suppressive Microenvironment Underpin Tumour Budding in Colorectal Cancer"

### *Supplementary Figure 8*

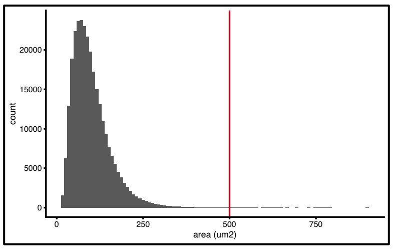


Segmentation & ReadCount & Visual Inspection
12 patients
210 FOVs
308520 cells
6519 genes

Cell Level QC
Single cell metrics: min counts per cell, negative probes
Spatial metrics: max area

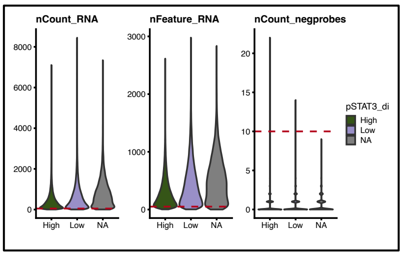


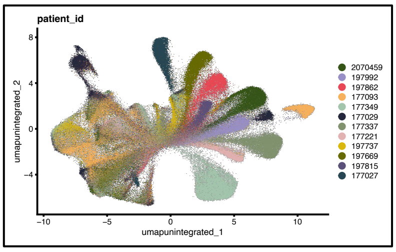


FOV Level QC
Diminished total counts
Biased gene expression profiles

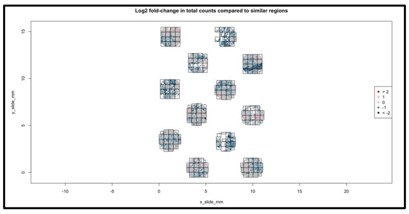


Normalisation
LogNorm

Batch effects
Harmony
scVI
fastMNN

| | Bio conservation | | | | | Batch correction | | | | | Aggregate score | | |
|---|---|---|---|---|---|---|---|---|---|---|---|---|---|
| Method | Isolated labels | KMeans NMI | KMeans ARI | Silhouette label | cLISI | Silhouette batch | iLISI | KBET | Graph connectivity | PCR comparison | Batch correction | Bio conservation | Total |
| X_pca_harmony | 0.48 | 0.00 | -0.00 | 0.51 | 0.41 | 0.94 | 0.32 | 0.17 | 0.74 | 0.96 | 0.63 | 0.28 | 0.42 |
| X_latent_scvi | 0.50 | 0.03 | 0.05 | 0.52 | 0.58 | 0.91 | 0.18 | 0.02 | 0.96 | 0.16 | 0.45 | 0.34 | 0.38 |
| X_pca_unintegrat | 0.49 | 0.00 | -0.00 | 0.51 | 0.84 | 0.89 | 0.05 | 0.02 | 0.86 | 0.08 | 0.38 | 0.37 | 0.37 |
